# Flexible computation of object motion and depth based on viewing geometry inferred from optic flow

**DOI:** 10.1101/2024.10.29.620928

**Authors:** Zhe-Xin Xu, Jiayi Pang, Akiyuki Anzai, Gregory C. DeAngelis

## Abstract

Vision is an active process. We move our eyes and head to acquire useful information and to track objects of interest. While these movements are essential for many behaviors, they greatly complicate the analysis of retinal image motion—the image motion of an object reflects both how that object moves in the world and how the eye moves relative to the scene. Our brain must account for the visual consequences of self-motion to accurately perceive the 3D layout and motion of objects in the scene. Traditionally, compensation for eye movements (e.g., smooth pursuit) has been modeled as a simple vector subtraction process. While these models are effective for pure eye rotations and 2D scenes, we show that they fail to apply to more natural viewing geometries involving combinations of eye rotation and translation. We develop theoretical predictions for how perception of object motion and depth should depend on the observer’s inferred viewing geometry. Through psychophysical experiments, we demonstrate novel perceptual biases that manifest when different viewing geometries are simulated by optic flow, in the absence of physical eye movements. Remarkably, these biases occur automatically, without training or feedback, and are well predicted by our theoretical framework. A neural network model trained to perform the same tasks exhibits neural response patterns similar to those observed in macaque area MT, suggesting a possible neural basis for these adaptive computations. Our findings demonstrate that the visual system automatically infers viewing geometry from optic flow and flexibly attributes components of image motion to either self-motion or depth structure according to the inferred geometry. Our findings unify previously separate bodies of work by showing that the visual consequences of self-motion play a crucial role in computing object motion and depth, thus enabling the visual system to adaptively perceive a dynamic 3D environment.

## 2 Introduction

From hawks catching prey to tennis players hitting a topspin forehand, humans and other animals frequently move their bodies to interact with the world. This requires processing sensory signals that arise from changes in the environment (e.g., objects moving in the world), as well as sensory signals that arise from our own actions (e.g., self-motion). A key challenge arises in these computations: the brain needs to decompose sensory signals into contributions caused by events in the environment and those resulting from one’s own actions. This type of computation is an example of causal inference (Kording et al., 2007; Shams and Beierholm, 2010; French and DeAngelis, 2020). To identify components of visual input that arise during self-motion, one must infer their own viewing geometry, namely, how the eyes translate and rotate relative to the scene as a result of eye, head, or body movements. As we will demonstrate, correctly computing the motion and 3D location of objects depends crucially upon correctly inferring one’s viewing geometry.

A classic example of how the brain compensates for visual consequences of action involves smooth pursuit eye movements, which we use to track objects of interest (Spering and Montagnini, 2011; Schütz et al., 2011). Many studies have examined how the brain compensates for the visual consequences of pursuit eye movements, and how this affects visual perception (Fleischl, 1882; Aubert, 1887; Filehne, 1922; Festinger et al., 1976; Wertheim, 1981, 1987; Swanston and Wade, 1988; Wertheim, 1994; Freeman and Banks, 1998; Freeman, 1999; Freeman et al., 2000; Haarmeier et al., 2001; Souman et al., 2005, 2006a; Spering and Gegenfurtner, 2007, 2008; Morvan and Wexler, 2009; Freeman et al., 2010; Schütz et al., 2011; Spering and Montagnini, 2011). For example, the Filehne illusion and Aubert-Fleischl phenomenon occur when the visual signal caused by a smooth eye movement is not accurately compensated (Fleischl, 1882; Aubert, 1887; Filehne, 1922). Theories that have been proposed to account for these perceptual phenomena (e.g., Mack and Herman, 1973; Festinger et al., 1976; Wertheim, 1981, 1987; Freeman and Banks, 1998; Freeman et al., 2010) generally share a common computational motif in which the brain compensates for the visual consequences of smooth pursuit by performing a vector subtraction of a reference signal that is related to eye velocity (Figure 1; Wertheim, 1987; Freeman and Banks, 1998; Spering and Gegenfurtner, 2008). While these models can account for biases in visual perception produced by a pure eye rotation, we demonstrate that they fail dramatically for even simple combinations of eye translation and rotation. Since the majority of previous empirical studies only tested visual stimuli on a 2D display, these limitations of the classic vector subtraction model appear to be largely unappreciated. In the present study, we show that computing object motion and depth in the world requires more than a simple vector subtraction, and that the brain flexibly interprets specific components of retinal image motion based on the 3D viewing geometry inferred from optic flow.

**Figure 1:**
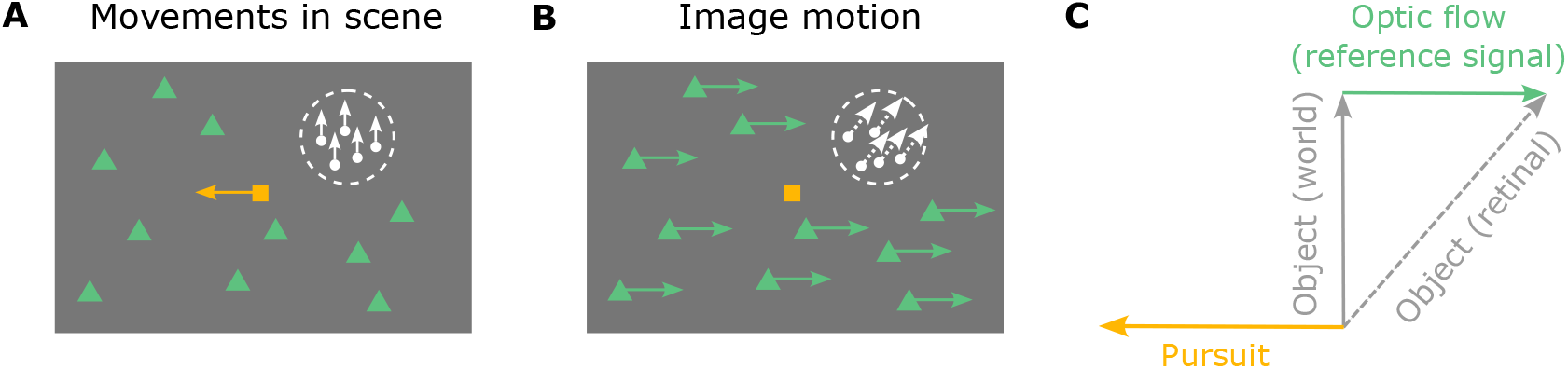
Schematic illustration of smooth pursuit eye movement and its visual consequences in a 2D scene. **A**, This diagram illustrates movements in the scene, including a pursuit target (yellow square) moving leftwards and an object (white patch of random dots) moving upwards. Green triangles depict stationary background elements in the scene. **B**, The resulting image motion for the scenario of panel A, shown in screen coordinates, assuming that the observer accurately pursues the yellow target. The image motion of the green triangles reflects optic flow generated by the eye movement (green arrows), and the image motion of the white object (white arrows) reflects both its motion in the world and the observer’s eye movement. **C**, The object’s motion in the world (solid gray arrow) can be obtained by subtracting the optic flow vector (green arrow) from the retinal image motion of the object (dashed gray arrow), which is equivalent to adding pursuit eye velocity (yellow arrow) to retinal image motion.

To illustrate the importance of 3D viewing geometry, consider how the effects of eye movement on visual input depend on scene structure. A typical paradigm for studying the effect of smooth pursuit on visual perception is shown in Figure 1A. A fixation target (yellow square) moves across the center of the screen, while a visual stimulus (e.g., a random-dot patch) appears at a particular location on the screen and moves independently (Figure 1A, white dots and arrows). The observer tracks the fixation target by making a leftward smooth pursuit eye movement, which results in rightward optic flow (Figure 1B, green arrows). For the moving object (white dots), retinal image motion reflects both its motion in the scene (world coordinates) and the optic flow resulting from eye movement. To compute its motion in the world, the observer needs to subtract the optic flow vector from the retinal image motion (or, equivalently, add eye velocity to it; Figure 1C). This scenario typically occurs when the observer remains stationary, and only the eyes rotate (Figure 2A). The object’s distance, or depth, does not affect the computation of its motion in world coordinates because the rotational flow field that results from a pure eye rotation is depth-invariant (Longuet-Higgins and Prazdny, 1980, Figure 2A). We hereby refer to this viewing geometry as Pure Rotation (R). See Video 1 for a demonstration of a pure rotational flow field.

**Figure 2:**
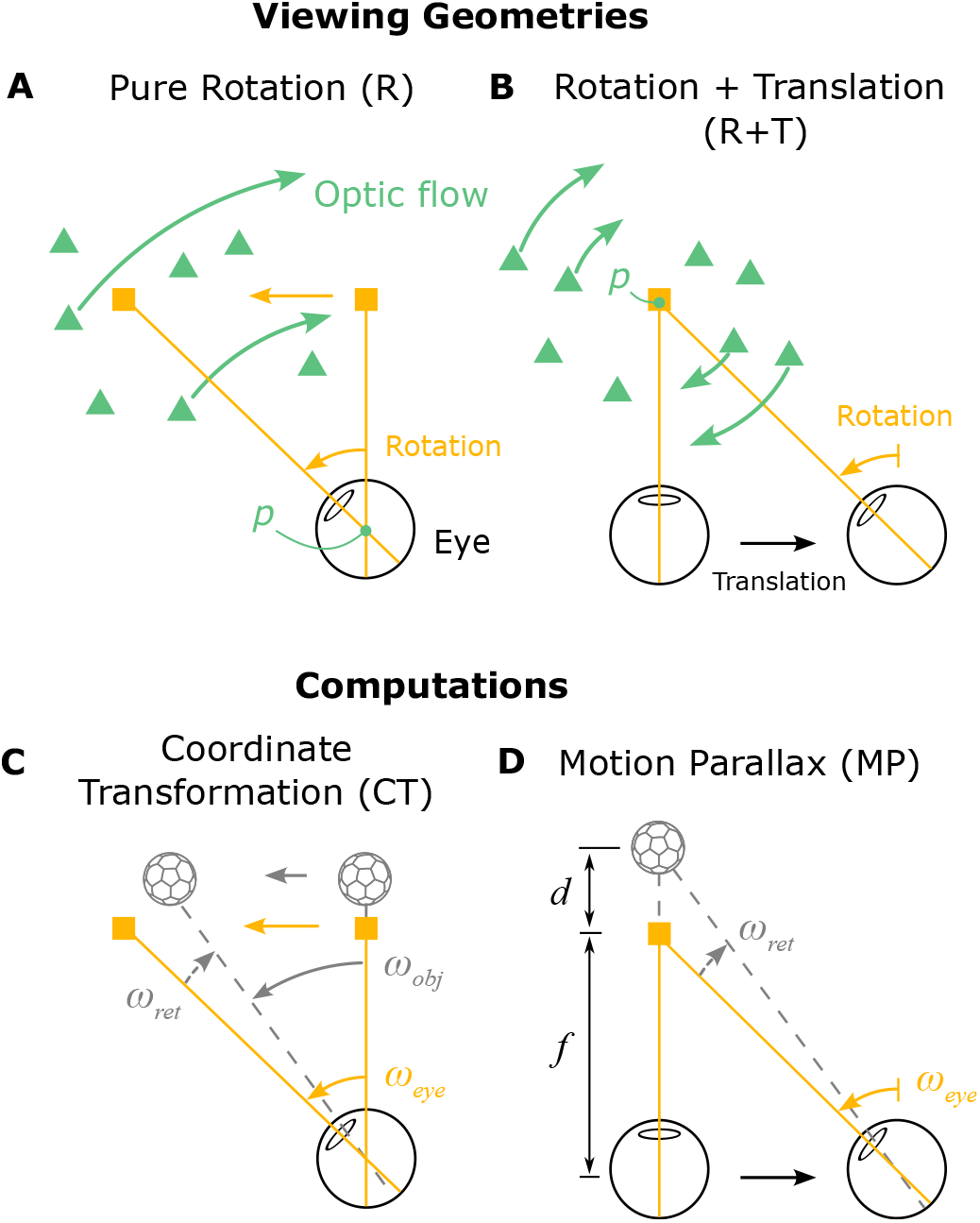
Schematic illustration of two viewing geometries and corresponding computations that can be performed. **A**, Top-down view of the Pure Rotation (R) viewing geometry, in which a stationary observer rotates their eye to track a moving fixation target (yellow square), resulting in an optic flow field (green arrows, shown for a subset of triangles for clarity) that rotates around the eye (rotation pivot, *p*) in the opposite direction of eye movement. **B**, In the Rotation + Translation (R+T) viewing geometry, the observer translates laterally and counter-rotates their eye to maintain fixation on a stationary target (yellow square), producing optic flow vectors (green arrows) in opposite directions for near and far objects (green triangles). This optic flow pattern is effectively a rotational flow field around the fixation target (rotation pivot, *p*). **C**, In the Pure Rotation (R) viewing geometry, the retinal image motion of a moving object (soccer ball shape), *ω*_*ret*_ (dashed gray arrow), reflects both its motion in the world, *ω*_*obj*_ (solid gray arrow), and the velocity of eye rotation, *ω*_*eye*_ (yellow arrow). By taking a vector sum between *ω*_*ret*_ and *ω*_*eye*_, the velocity of the object can be transformed from retinal coordinates to world coordinates, hereafter referred to as a coordinate transformation (CT). **D**, In the Rotation + Translation (R+T) viewing geometry, the retinal image motion of a stationary object (soccer ball shape), *ω*_*ret*_ (dashed gray arrow), depends on where the object is located in depth, *d*, and the rotational eye velocity, *ω*_*eye*_ (yellow arrow). By computing the ratio between *ω*_*ret*_ and *ω*_*eye*_, the depth of the object can be obtained from the motion parallax (MP) cue.

Now consider a simple extension of the viewing geometry in which the observer translates laterally while maintaining visual fixation on a fixed point in the scene by counter-rotating their eyes (Figure 2B). We refer to this viewing geometry as Rotation + Translation (R+T). By introducing this lateral translation, the same leftward pursuit eye movement becomes associated with a drastically different optic flow pattern (Figure 2B, green arrows). In this geometry, the scene rotates around the point of fixation and motion parallax (MP) cues for depth become available.

Stationary objects at different depths relative to the fixation point move with different velocities on the retina, and the retinal image speed increases with distance from the fixation point (Figure 2B; see Video 2 for a demonstration). Stationary elements in the scene nearer than the fixation point move in the same direction as the eye, while far elements move in the opposite direction (Figure 2B, green triangles and arrows). Because there are a variety of optic flow vectors associated with the same eye movement in the R+T geometry, compensating for the visual consequences of pursuit can no longer be a simple vector subtraction; one must consider the depth of objects.

Therefore, the visual consequences of a smooth pursuit eye movement differ greatly depending on viewing geometry, and it is crucial to understand that different computations are typically performed to interpret the scene in these two viewing geometries: coordinate transformation in the R geometry and estimation of depth from motion parallax in the R+T geometry (Figure 2C and D). Coordinate transformation (CT) refers to transforming object motion from retina-centered coordinates to world-centered coordinates (Figure 2C; e.g., Andersen et al., 1993; Freeman and Banks, 1998; Swanston et al., 1992; Wade and Swanston, 1996). In the R geometry, since it is extremely unlikely that the entire scene rotates around the eye due to external causes, it is natural for the brain to attribute rotational optic flow to eye rotation and to represent object motion relative to the head by subtracting optic flow from the retinal image. For example, leftward smooth pursuit would induce rightward optic flow and a horizontal (leftward) bias in perceived direction of a moving object relative to its image motion, as observed empirically (Souman et al., 2005; Zivotofsky et al., 2005; Champion and Freeman, 2010). On the other hand, in the R+T geometry, motion parallax (MP) provides valuable information about the depth of stationary objects (e.g., Rogers and Rogers, 1992; Nawrot, 2003; Naji and Freeman, 2004; Nawrot and Stroyan, 2009, 2012). Specifically, depth can be computed as the ratio of its retinal image motion and the pursuit eye velocity (Figure 2D; Nawrot and Stroyan, 2009). When the eye translates and counter-rotates horizontally, as illustrated in Figure 2B, the horizontal component of an object’s motion could be attributed to depth, especially if other depth cues are not in conflict. Therefore, in the R+T geometry, it is natural for the brain to attribute at least some of the horizontal component of motion to depth while the vertical component is attributed to independent object motion, thus leading to a vertical bias in perceived direction of the object. Thus, as we formalize below, accounting for the visual consequences of eye movements under different viewing geometries predicts systematic patterns of biases in motion and depth perception that otherwise may not be anticipated.

CT computations and estimation of depth from MP have been studied extensively in terms of behavior (e.g., de Graaf and Wertheim, 1988; Filehne, 1922; Wertheim, 1987; Freeman and Banks, 1998; Mack and Herman, 1973; Nawrot, 2003; Naji and Freeman, 2004; Nawrot et al., 2014; Nadler et al., 2009; Niehorster and Li, 2017; Ono et al., 1986; Rogers, 1993; Rogers and Graham, 1979; Rushton and Warren, 2005; Swanston et al., 1992; Wade and Swanston, 1996; Wallach et al., 1985; Warren and Rushton, 2009; Wertheim, 1987) and neural mechanisms (e.g., Brostek et al., 2015; Chukoskie and Movshon, 2009; Ilg et al., 2004; Inaba et al., 2007, 2011; Kim et al., 2015, 2022; Nadler et al., 2008, 2009; Sasaki et al., 2020; Thier and Erickson, 1992; Xu and DeAngelis, 2022). However, previous studies generally treat these two phenomena as separate and unrelated. Interestingly, in these two viewing geometries, the same two signals—retinal velocity and eye velocity—are typically combined in different ways: summation to compute object motion in the world (CT, Equation 3), and division to estimate depth based on MP (Equation 4). This raises an important question: how does the brain infer the relevant viewing geometry and use this information to compute object motion and depth in a context-dependent fashion?

We demonstrate that the R and R+T viewing geometries are specific instances of a general framework that explains interactions between motion and depth perception under a range of self-motion conditions. While the computations for object motion and depth take apparently distinct forms, involving addition and division respectively, they can be unified under the same framework when considering viewing geometry. We conduct psychophysical experiments to characterize the effects of viewing geometry, simulated by optic flow, on the perception of object motion and depth. Our findings show that humans flexibly and automatically (without any training or feedback) compute motion and depth based on the simulated viewing geometry, even in the absence of extra-retinal signals about eye movement. Rather than being a nuisance variable to suppress, visual image motion induced by smooth eye movements provides a powerful input for flexibly computing object motion and depth in a context-specific manner.

To investigate potential neural substrates of these flexible computations of motion and depth, we train recurrent neural networks to perform these tasks and compare the representations learned by hidden units of the network with those in the primate visual cortex. We show that task-optimized recurrent neural networks exhibit adaptive representations roughly similar to those found in neurons in the macaque middle temporal (MT) area. Our work thus reveals the computational principles and a potential neural basis of how we perceive the 3D world while in motion.

## 3 Results

We present a novel theory that links computations of object motion and depth with viewing geometry by considering the optic flow patterns produced by eye rotation and translation. Our theory predicts striking differences in motion and depth perception between the R and R+T viewing geometries, which we validate by performing psychophysical experiments with human subjects. We demonstrate that humans automatically and flexibly alter their estimation of motion and depth based on the viewing geometry simulated by optic flow. Finally, we show that recurrent neural networks trained to compute object motion and depth exhibit non-separable retinal and eye velocity tuning similar to neurons found in area MT, suggesting a potential neural basis for computing motion and depth in a viewing geometry–dependent manner.

### 3.1 Different viewing geometries generate distinct optic flow fields

We start by asking which cues are pivotal in shaping beliefs about viewing geometry. While the retinal image motion of the object and eye-in-head rotation may be identical between the R and R+T viewing geometries (Figure 2C and D), the optic flow field generated by eye movements clearly differentiates the viewing geometry (Figure 2A and B; Videos 1 and 2). In the R geometry, the observer’s head is stationary, and only the eyes rotate; therefore, the optic flow field reflects rotation of the scene around the eye (Figure 2A, point *p*). In the R+T geometry, the observer’s head translates laterally, and the eyes counter-rotate to maintain fixation on a world-fixed point. Therefore, the optic flow field reflects rotation of the scene around the fixation point (Figure 2B, point *p*).

In essence, the main difference between R and R+T viewing geometries can be captured by a single parameter: the rotation pivot of the optic flow field produced by eye movements. To formalize the relationship between retinal image motion, eye rotation, object motion, and depth, we consider a more general viewing geometry depicted in Figure 3; this generalization encompasses the R and R+T geometries. The retinal image motion of an object, *ω*_*ret*_, is a combination of the object’s scene-relative motion, *ω*_*obj*_, motion parallax from the observer’s translation (which depends on depth), 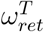, and optic flow produced by the observer’s pursuit eye movement, 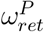 (Figure 3B):

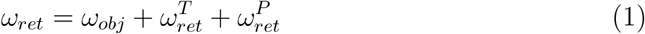

**Figure 3:**
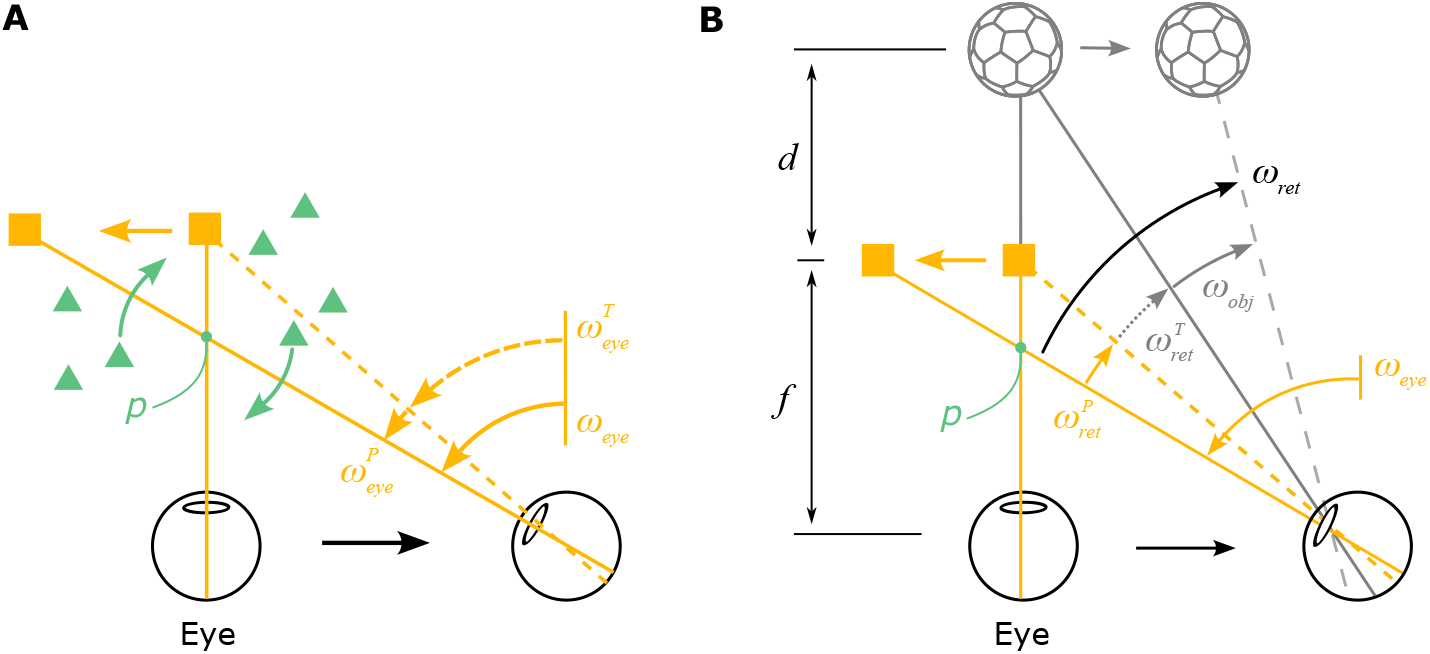
Geometry of a more generalized viewing scenario. **A**, The eye translates to the right while maintaining fixation on a moving target (yellow square) by making a smooth eye movement with velocity *ω*_*eye*_. The eye velocity is comprised of a compensatory rotation for the translation, 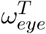, and a component related to tracking the moving target, 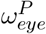 . The rotation pivot of the optic flow field, *p*, is located between the eye and the fixation target. The amplitude of eye translation and rotation is exaggerated for the purpose of illustration. **B**, When an object (soccer ball shape) is located at depth, *d*, and moves independently in the world, its retinal image velocity, *ω*_*ret*_, is determined by its own motion in the world, *ω*_*obj*_, motion parallax produced by the observer’s translation, 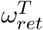, and image motion resulting from the pursuit eye movement, 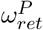 .

Therefore, object motion in world coordinates can be expressed as (see Supplementary Information for derivation):

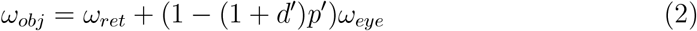

Here, *ω*_*obj*_, *ω*_*ret*_, and *ω*_*eye*_ denote the angular velocities of object motion in world coordinates, its retinal motion, and eye rotation. *d*^*′*^ represents the object’s depth, *d*, normalized by viewing distance, *f* : *d*^*′*^ ≜ *d/f* . When *d*^*′*^ = 0, the object is at the same depth as the fixation plane, whereas *d*^*′*^ *<* 0 means near and *d*^*′*^ *>* 0 means far compared to the fixation plane. Similarly, *p*^*′*^ represents the normalized rotation pivot, *p*^*′*^ ≜ *p/f*, where *p* is the distance from the rotation pivot to the cyclopean eye. Therefore, *p*^*′*^ = 0 corresponds to the R geometry and *p*^*′*^ = 1 indicates the R+T geometry.

When *p*^*′*^ = 0 (R geometry), object motion in world coordinates is the sum of retinal and eye velocities, thus capturing the coordinate transformation computation:

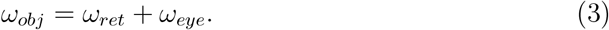

When *p*^*′*^ = 1 (R+T geometry) and the object is stationary in the world, *ω*_*obj*_ = 0, the object’s relative depth is the ratio between retinal and eye velocities, resulting in the approximate form of the motion-pursuit law (Nawrot and Stroyan, 2009):

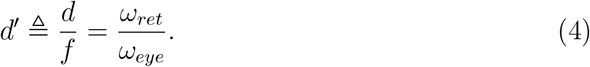

Our derivation of Equation (2) (see Supplementary Information for details) thus provides a general framework that includes the R and R+T viewing geometries, and thereby links together the computations of object motion and depth for a moving observer. While the addition and division computations for computing motion and depth appear quite different on the surface, both operations can be expressed as a single computation when we incorporate the rotation pivot of optic flow. Thus, it suggests that the brain transitions between these operations when optic flow implies different viewing geometries.

This finding raises important questions. Do people use optic flow to infer their viewing geometry? Do they perceive object motion and depth differently when viewing optic flow that simulates different viewing geometries? We conducted a series of psychophysical experiments to measure motion and depth perception in humans while presenting optic flow patterns simulating different viewing geometries. We also examined whether effects are substantially different when subjects pursue a target with their eyes, as compared to when pursuit is visually simulated.

### 3.2 Humans rely on 3D viewing geometry to infer motion in the world

In Experiment 1, human participants performed a motion estimation task. Specifically, we presented an object moving with a fixed set of directions on the retina while simulating different viewing geometries with large-field background motion (Figure 4; see Methods for details). The object and optic flow were presented for 1 s, after which a probe stimulus appeared, and participants adjusted a dial to match the motion direction of the probe with that of the object (Figure 6A and B).

**Figure 4:**
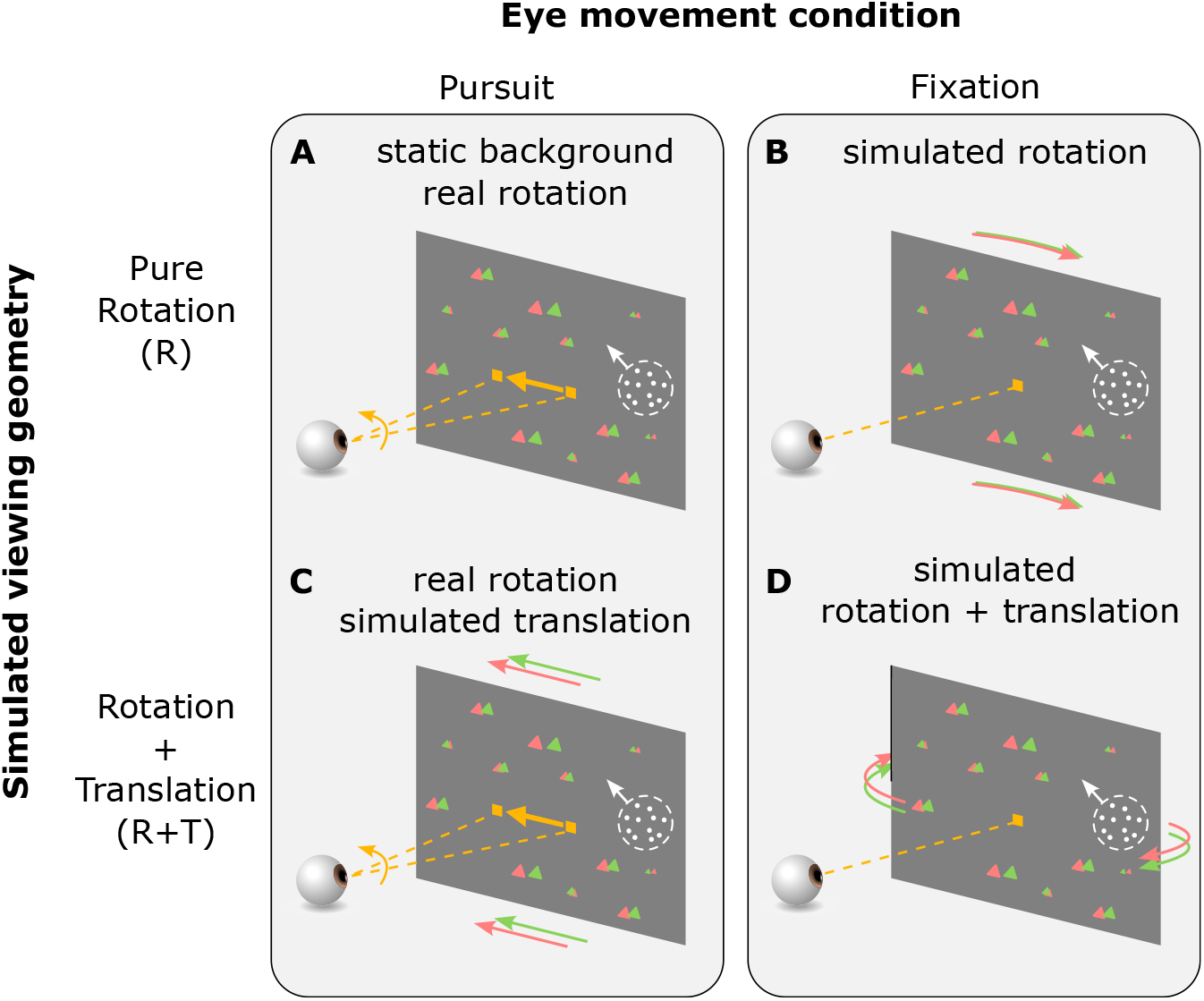
Stimulus and task conditions for Experiment 1. In the Pursuit conditions (**A & C**), the fixation target moved horizontally across the screen during the stimulus presentation period. Participants tracked the target by making smooth pursuit eye movements. In the Fixation conditions (**B & D**), the fixation target remained stationary at the center of the screen and participants maintained fixation on the target throughout the trial. In the R viewing geometry (**A & B**), a pure eye rotation was either executed by the participant in the Pursuit condition (**A**, yellow arrow) or simulated by background optic flow (**B**, red/green triangles). In the R+T viewing geometry (**C & D**), lateral translation of the eye relative to the scene was always simulated by background optic flow, and eye rotation was either real or simulated, as in the R geometry.

Four main experimental conditions were interleaved: two eye-movement conditions times two simulated viewing geometries (Figure 4; Videos 3–6). The two eye-movement conditions include: (1) the Pursuit condition, in which participants tracked a moving target by making smooth pursuit eye movements while the head remained stationary (Figure 4A and C), and (2) the Fixation condition, in which participants fixated on a stationary target at the center of the screen, and eye movements were simulated by background motion (Figure 4B and D). The two background conditions include: (1) the R condition, in which background dots simulated the R viewing geometry (Figure 4A and B), and (2) the R+T condition, in which the background simulated the R+T geometry (Figure 4C and D). Simulated eye translation in the R+T condition was always horizontal (i.e., along the interaural axis), and target motion in the Pursuit condition was also always horizontal. Thus, all real and simulated pursuit eye movements were horizontal (leftward or rightward). Notably, participants did not receive any form of feedback during performance of the task. In addition to the four main conditions, a control condition, in which no background dots were present and thus no cues for viewing geometry were available, was interleaved to measure baseline motion estimation performance (Videos 7–8).

Because the eye translation and/or rotation simulated by background motion was always horizontal, we expect the two viewing geometries, R and R+T, to affect perception of the object’s horizontal motion component. For the R viewing geometry, in which the brain naturally performs CT, horizontal eye velocity should be added to the object’s retinal motion (Equation 3), resulting in a perceptual bias towards the horizontal (Figure 5, top row, orange arrows; Figure 6C, orange curves). Because the object was presented monocularly and its size was kept constant on the screen across conditions, the object’s depth should be ambiguous in the R geometry (Figure 5, top row, purple; Figure 7C, orange band), for which optic flow is depth-invariant. Conversely, in the R+T case, we expect eye velocity to be combined with the object’s horizontal retinal motion to compute depth from MP based on the motion-pursuit law (Equation 4; Nawrot and Stroyan, 2009). Because of the absence of other depth cues, we hypothesize that the horizontal component of the object’s retinal motion will be explained away as MP resulting from observer translation (Figure 5, bottom row, purple arrows). Consequently, only the remaining vertical motion component will be perceived as object motion (Figure 5, bottom row, blue arrows). In this case, we expect participants to show a perceptual bias toward vertical directions (Figure 5, bottom row, blue arrows; Figure 6C, blue curves).

**Figure 5:**
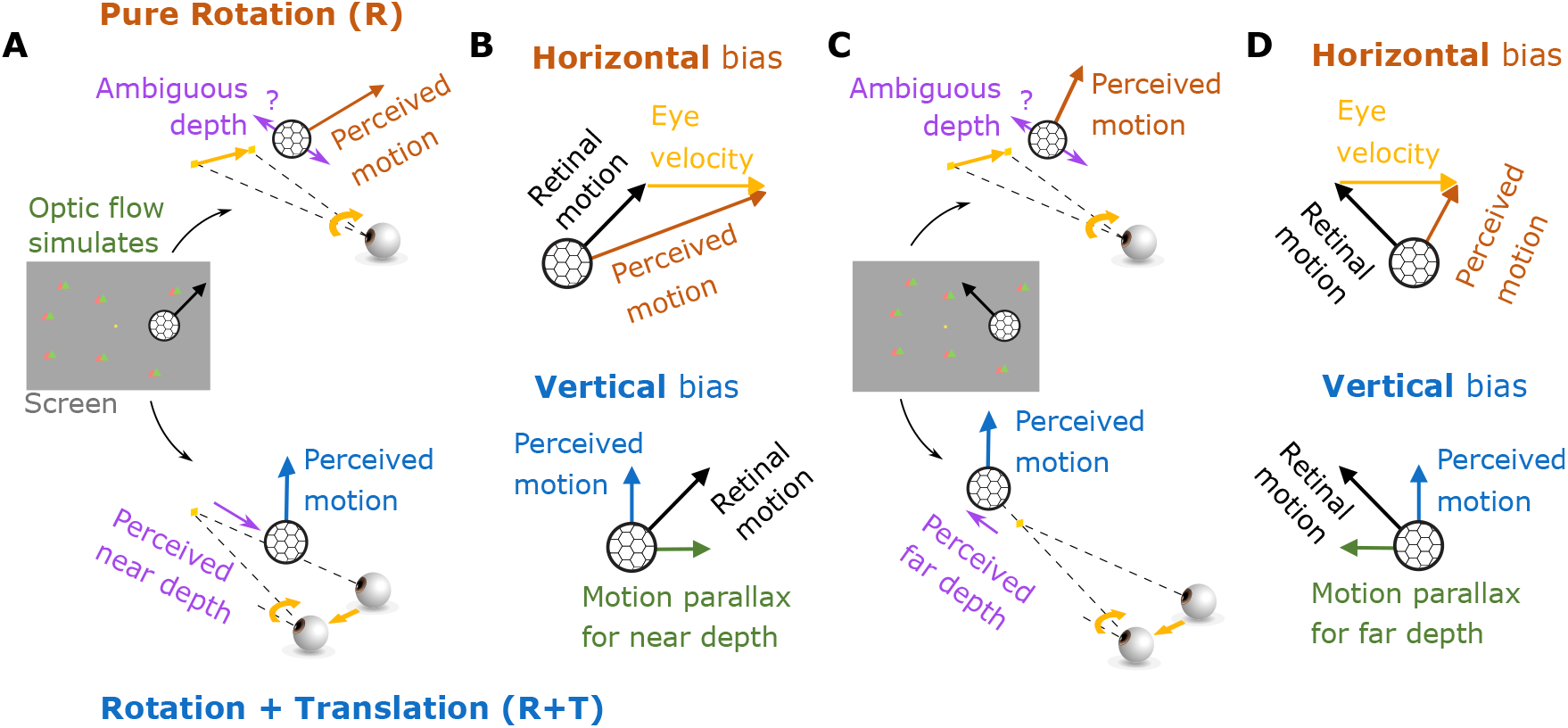
Predictions for motion and depth perception in the R and R+T viewing geometries. **A**, Center, stimulus display showing a fixation target (yellow square) and an object (soccer ball shape) moving up and to the right (illustration of the Fixation condition). Meanwhile, background dots (red/green triangles) simulate the R (top) or R+T (bottom) viewing geometries. Top, in the R geometry, a rightward eye velocity (yellow arrow) is added to the image motion of the object, resulting in a rightward bias in motion perception (orange arrow). The object’s depth remains ambiguous due to the absence of reliable depth cues. Bottom, in the R+T geometry, the horizontal component of the object’s image motion can be explained away as motion parallax, such that the object is perceived as having a near depth. The residual vertical motion is then perceived as the object’s motion in the world (blue arrow), leading to a vertical bias in perception. **B**, Summary of the relationships between retinal motion, eye velocity, and perceived object motion in the R (top) and R+T (bottom) geometries. **C**, Same format as **A**, except that the object moves up and to the left on the display (center panel). In the R geometry (top), depth remains ambiguous and the same rightward eye velocity is added to the object’s motion, again producing a rightward bias. In the R+T geometry (bottom), the horizontal component of the object’s image motion reverses direction, thus causing a far-depth percept. The perceived motion remains vertical. **D**, Same format as **B**, except for the scenarios depicted in **C**.

**Figure 6:**
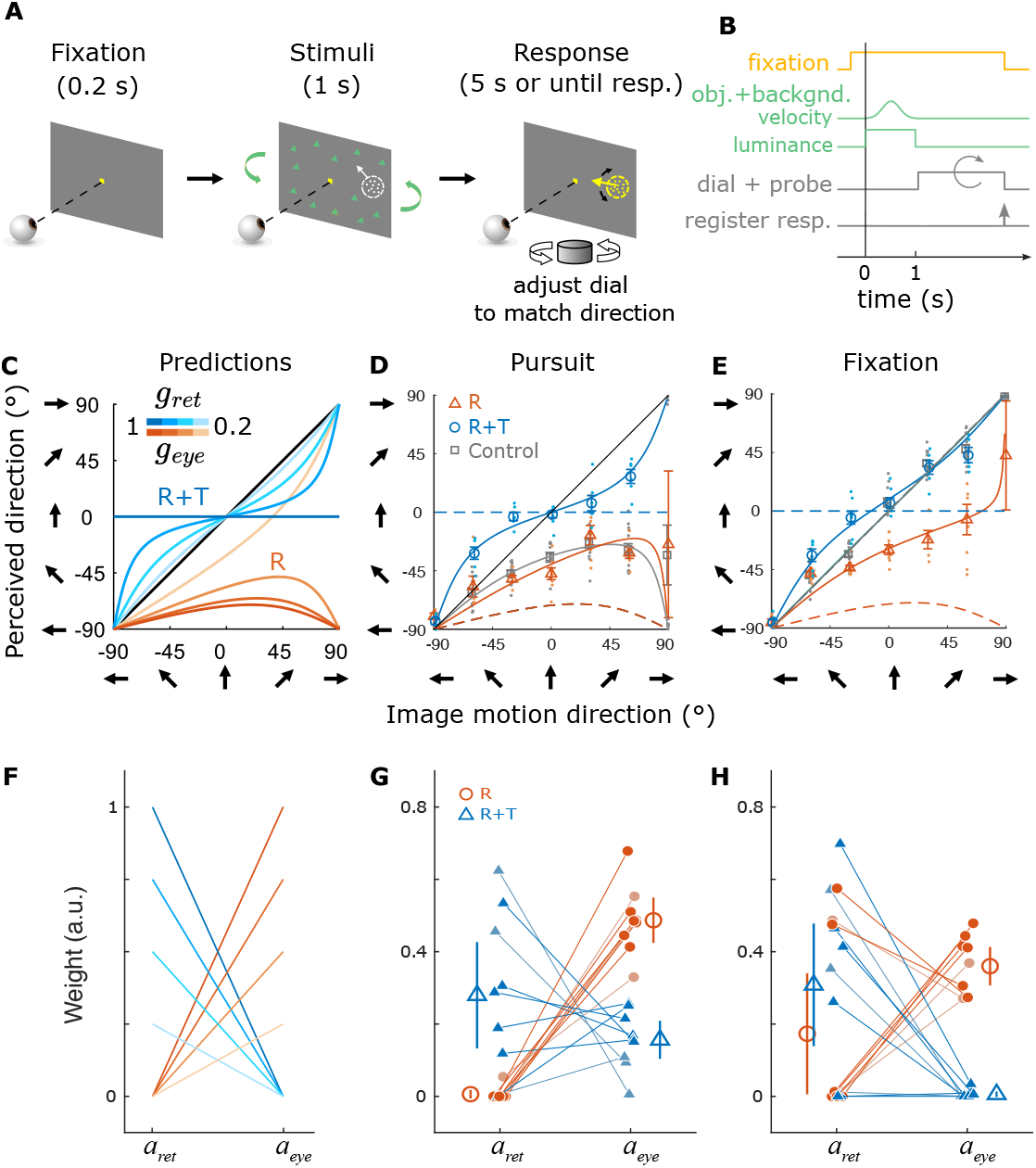
Procedure, predictions, and results for the motion estimation task. **A**, At the beginning of each trial, a fixation target (yellow square) was presented at the center of the screen. After fixation, the visual stimulus appeared, including the background dots (green triangles) and the moving object (white dots). The 1-s stimulus presentation was followed by a response period, during which a probe stimulus composed of random dots (yellow dots) appeared, and the subject turned a dial to match the probe’s motion direction with the perceived direction of the object. **B**, Time course of stimulus and task events for the motion estimation task. **C**, Prediction of perceived motion direction in the R (orange curves) and R+T (blue curves) geometries. In the R+T geometry, we expect a bias toward the vertical direction (0^°^on the y-axis), whereas a horizontal bias (toward -90^°^on the y-axis) is predicted in the R geometry. The color saturation of the curves depends on the gains in Equation 9 (*g*_*ret*_ and *g*_*eye*_, respectively). **D–E**, Data from one example subject (h500) in the Pursuit (**D**) and the Fixation (**E**) conditions. Individual dots represent the reported direction in each trial, and open markers indicate averages. Dashed curves are the predictions of the R (orange) and R+T (blue) geometries with *g*_*ret*_ = *g*_*eye*_ = 1, and solid curves are linear model fits to the data. Error bars indicate 1 SD. **F**, Predictions of the weights for retinal and eye velocities in the R (orange) and R+T (blue) geometries. Shading indicates the gains as in **C. G**, Weights of retinal and eye velocities in the pursuit condition. Each filled circle and triangle represents data from one participant, and open symbols indicate means across participants. Error bars show 95% CIs. **H**, Weights in the fixation condition. Format as in **G**.

**Figure 7:**
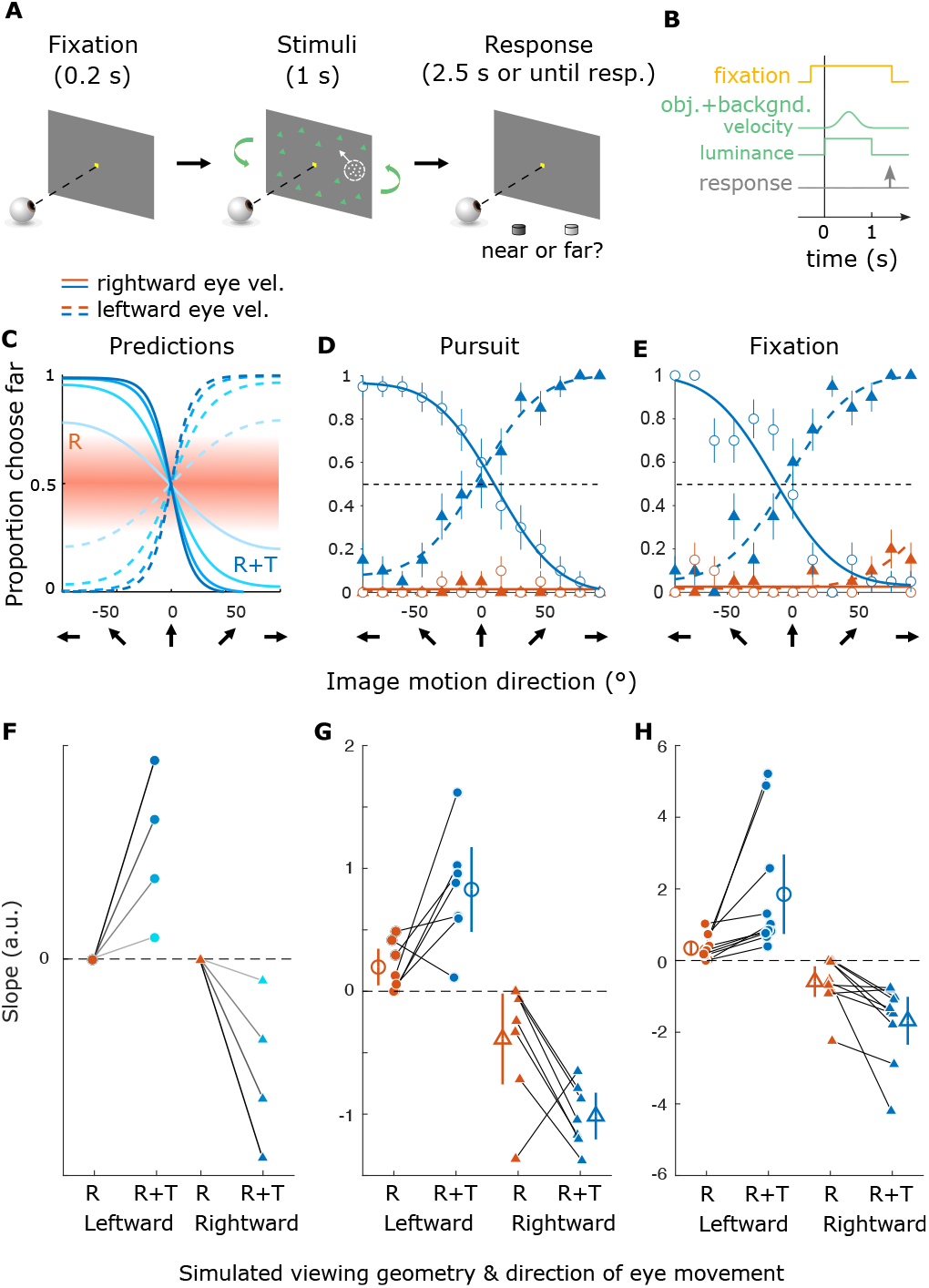
Procedure, predictions, and results for the depth discrimination task. **A**, In each trial, a fixation point was followed by presentation of the visual stimuli, including the object and background. During or after stimulus presentation, participants pressed one of two buttons to report whether the object was perceived to be near or far relative to the fixation point. **B**, Time course of stimulus and task events for the depth discrimination task. **C**, Model predictions for depth perception in the R+T (blue curves) and R (orange) viewing geometries. Dashed and solid curves indicate leftward and rightward (real or simulated) eye rotations, respectively. Color saturation of the blue curves indicates the amount of eye movement accounted for in the prediction (same as Figure 6C). Orange band represents the ambiguity of depth in the R viewing geometry. **D–E**, Psychometric curves from a naïve subject (h507) in the Pursuit (**D**) and Fixation (**E**) conditions. Error bars indicate S.E.M. **F**, Predicted slopes of psychometric functions in each simulated viewing geometry. Circles and triangles denote slopes for leftward and rightward eye movements, respectively. Orange and blue symbols represent the R and R+T viewing geometries, respectively. Saturation of symbol color indicates the gains in Equation (10), same as in **C. G**, Slopes of psychometric functions in the Pursuit conditions for all subjects. Each solid symbol represents one participant, and large open markers show the group average. Error bars indicate 95% CIs. **H**, Slopes of psychometric functions in the Fixation conditions. Format as in **G**.

Observers may not accurately infer the viewing geometry from optic flow. In the R geometry, they might underestimate their eye velocity; in the R+T geometry, they might only attribute a portion of the horizontal velocity component to depth. Therefore, our theoretical framework incorporates two discount factors into our predictions (Equation 9; Figure 6C), which are conceptually analogous to a flow parsing gain (Niehorster and Li, 2017; Peltier et al., 2020, 2024).

Indeed, we found systematic biases in motion direction reports for all participants, and the pattern of biases is generally consistent with our predictions (compare Figure 6D–E and Figure S1 with Figure 6C). Specifically, we found an overall bias toward reporting horizontal directions in the R viewing geometry for both real (Pursuit) and simulated eye rotation (Fixation) conditions (Figure 6D–E and Figure S1, orange).

In the R+T viewing geometry, we found an overall bias toward reporting vertical directions in the Fixation condition, with mixed results for the Pursuit condition as described further below (Figure 6D–E and Figure S1, blue). As a control, we found no bias in the Fixation condition when there were no background dots, indicating that participants accurately perceived image motion on the screen when there was no background motion or pursuit eye movements (Figure 6E and Figure S1, gray dots and squares). In the Pursuit condition with no background dots, participants showed a horizontal perceptual bias, suggesting a partial coordinate transform toward world coordinates (Figure 6D, gray). This result is consistent with previous findings that motor commands associated with real pursuit modulate motion perception (e.g., Wertheim, 1987; Freeman and Banks, 1998; Spering and Gegenfurtner, 2008). For some subjects, the overall response pattern in the R+T Pursuit condition deviated substantially from the identity line and shifted toward the lower half of the plot (Figure S1A, C, D, I–L). This might indicate that these subjects interpreted the viewing geometry as a mixture between R and R+T (see Discussion).

Because Equation (2) shows that perceived object motion can be expressed as a linear combination of retinal and eye velocities in both R and R+T geometries, we fit a linear model to each participant’s responses in these experimental conditions. The linear model incorporated two parameters: a weight *a*_*ret*_, which scales down the horizontal component of retinal velocity, and a weight *a*_*eye*_, which accounts for the addition of eye velocity to retinal velocity. If *a*_*ret*_ = 0, the horizontal component of retinal velocity is attributed to object motion (Figure 6F, orange). In contrast, if *a*_*ret*_ *>* 0, this suggests that part of the horizontal component is interpreted as MP (Figure 6F, blue). The parameter *a*_*eye*_ determines the extent to which eye velocity is added to the retinal velocity (Equation 10; see Methods for details). Overall, this simple linear model nicely captured the response patterns across individual subjects (Figure 6D and E, Figure S1; solid curves).

Equation (2) predicts that in the R geometry, *a*_*ret*_ should be close to zero because retinal motion of the object can be fully explained as object motion in the world, and *a*_*eye*_ should be greater than zero because eye velocity is added to retinal velocity to achieve CT (Figure 6F, orange lines). In the R+T geometry, because at least a portion of the horizontal retinal motion can be explained away as depth from MP, *a*_*ret*_ should be greater than zero such that only part of the horizontal retinal velocity is perceived as object motion, whereas *a*_*eye*_ should be close to zero because eye velocity would not be added to the object’s motion (Figure 6F, blue lines).

For most subjects, the estimated weights align reasonably well with these predictions (Figure 6G and H). We found greater values of *a*_*ret*_ in the R+T condition than the R condition, and greater values of *a*_*eye*_ in the R condition than the R+T case. This pattern is consistent across the majority of the participants and between the pursuit and fixation conditions (*p <* .001 for 8 out of 9 participants, 88.9%, in the Pursuit condition and for 5 out of 9 participants, 55.6%, in the Fixation conditions, Wilcoxon signed-rank test on 500 bootstrapped resamples with replacement for each participant). At the group level, *a*_*ret*_ had a significantly greater value in R+T compared to the R condition during pursuit (∆median = 0.313, *p* = 7.813 × 10^*−*3^, Wilcoxon signed-rank test) but not during fixation (∆median = 0.269, *p* = 0.496, Wilcoxon signed-rank test); *a*_*eye*_ had a significantly greater value in the R condition compared to R+T condition during both pursuit (∆median = 0.336, *p* = 3.91 × 10^*−*3^, Wilcoxon signed-rank test) and fixation (∆median = 0.370, *p* = 3.91 × 10^*−*3^, Wilcoxon signed-rank test). It is worth noting that there is considerable variability across individuals, most notably three participants who are outliers in the R geometry during fixation (Figure 6H, orange lines). This variability might be due to participants’ varying ability to infer viewing geometry from optic flow, biases in estimating eye velocity, and/or variability in pursuit execution.

As noted above, in the R+T viewing geometry, many participants showed an overall bias towards the lower half of the plot in the Pursuit condition (Figure S1A, C, D, I–L, blue). Although this pattern deviates from our predictions (Figure 6C), it is effectively captured by values of *a*_*eye*_ (Figure 6G, blue) that are greater than zero in our linear model fit for the Pursuit condition, which may suggest that some subjects still perform a partial CT in the R+T geometry.

### 3.3 Viewing geometry biases depth perception based on motion parallax

The findings of the previous section demonstrate that motion perception of most subjects is systematically biased by viewing geometry, in agreement with our theoretical predictions. Our analysis of viewing geometry, as described by Equation (2), also makes specific predictions for how depth perception should vary between the R and R+T viewing geometries. In the case of R, the optic flow is depth invariant, the object was viewed monocularly, and the size of dots was kept constant across conditions. Therefore, there was no information available to form a coherent depth percept. When asking participants to judge whether the object was near or far compared to the fixation plane, we expect the response to be at chance or biased to an arbitrary depth sign based on their prior beliefs (Figure 5A and C, top-right, purple; Figure 7C, orange band). In the R+T geometry, we expect the horizontal component of the object’s retinal motion to be explained away as MP (Figure 5A and C, bottom-right, purple). Therefore, the ratio between the horizontal component of retinal motion and the simulated eye velocity should determine the object’s depth, based on the motion-pursuit law (Nawrot and Stroyan, 2009). As retinal direction of the object changes, the horizontal component of its retinal motion varies from negative to positive, yielding a change in perceived depth from near to far, or vice-versa. When asked to judge the depth sign (i.e., near or far), we expect participants to show inverted psychometric curves for opposite directions of eye movement, since perceived depth sign depends on both the sign of retinal velocity and the sign of eye velocity (Figure 7C, blue curves).

In Experiment 2, we tested these predictions for perceived depth by asking human participants to discriminate the object’s depth in the two viewing geometries (Figure 7A and B). The stimuli were the same as in Experiment 1, except for the following: 1) the retinal direction of the object ranged from 0^°^ to 180^°^ instead of 0^°^to 360^°^ and 2) after stimulus presentation, subjects reported the perceived depth of the object (near or far relative to the fixation point) by pressing one of two buttons corresponding to each percept. Figure 7D and E show the results from one participant. In the R condition, the participant performed poorly, almost always reporting the object to be near, and no systematic difference was found between the two directions of simulated eye movement (Figure 7D–E, orange). Conversely, in the R+T condition, we observed a clear transition in depth reports from near to far, or vice-versa, as a function of retinal motion direction, and the psychometric curve inverted when the direction of eye movement was reversed (Figure 7D-E, blue), consistent with our predictions (Figure 7C).

Figure S2 shows similar data from other participants. Some subjects show substantial non-zero slopes in the R condition, but almost always with lower magnitudes than in the R+T condition. Because the two viewing geometries were randomly interleaved within each session, we speculate that some subjects might learn the inherent association between eye movement direction, retinal motion, and depth sign in the R+T viewing geometry and might generalize this association to the R viewing geometry (e.g. Figure S2G). Critically, because the retinal image motion of the object was identical between the R and R+T conditions, differences in depth perception can only be explained by the difference in optic flow patterns between the two viewing geometries.

The observed patterns of results are broadly consistent across most participants, as summarized in Figure 7F–H. We observed a significantly greater magnitude of slope in the R+T geometry compared to the R geometry, for 6 out of 7 (85.7%) participants in the Pursuit condition and 8 out of 10 (80%) participants in the Fixation condition (*p* < 0.05, permutation test for each participant). At the group level, the magnitude of the slope of the psychometric function is greater in the R+T geometry compared to the R geometry, indicating a stronger depth percept (∆median |slope| = 0.800, *p* = 3.05 × 10^*−*3^ for the Pursuit condition; ∆median |slope| = 0.693, *p* = 1.20 × 10^*−*4^ for the Fixation condition; Wilcoxon signed-rank test across participants). The distinct results between the R and R+T viewing geometries suggest that the main contribution of optic flow to depth perception observed here is generated by the combination of translation and rotation of the eye, which produces optic flow with a rotation pivot point at the fixation target. These results also provide the first direct evidence that humans automatically perceive depth from MP when eye rotation is inferred from optic flow, as proposed by Kim et al. (2015) (also see Buckthought et al., 2017).

Together, the results of Experiments 1 and 2 suggest that human subjects automatically, and without any training, infer their viewing geometry from optic flow and subsequently perform the more natural computation in each geometry (CT for the R geometry, and depth from MP for the R+T geometry). As a result, the interaction between retinal and eye velocity signals automatically switches from summation (for CT) to division (for depth from MP) based on the inferred viewing geometry.

### 3.4 Underlying neural basis implied by task-optimized recurrent neural network

So far, we have demonstrated that the visual system flexibly computes motion and depth based on optic flow cues to viewing geometry. How does the brain adaptively implement these computations? Recurrent neural networks (RNNs) have proven to be a useful tool for answering this type of question (Mante et al., 2013; Pandarinath et al., 2018; Rajan et al., 2016; Yang et al., 2019). RNNs trained on specific tasks could perform these tasks with precision comparable to humans and showed response dynamics resembling those observed in biological systems (e.g., Mante et al., 2013; Pandarinath et al., 2018; Rajan et al., 2016; Yang et al., 2019).

We trained an RNN with 64 recurrent units to perform the motion estimation and depth discrimination tasks given retinal image motion of a target object and different optic flow patterns (Figure 8A). At each time point, recurrent units in the network receive three inputs: retinal motion of the object and optic flow vectors of two background dots located at different depths. Because only horizontal rotation and translation of the eye were tested in our psychophysical experiments, the inputs to the network only included the horizontal velocity of the object’s retinal motion and two optic flow vectors. Two scalar outputs are produced: the horizontal velocity of world-centered object motion and depth relative to the fixation point. We chose these inputs and outputs to approximate the structure of the psychophysical task while keeping the network architecture simple. Because of the layout of the rotational pivot in the R and R+T geometries (Figure 2A and B), the optic flow vectors of two background dots, one at a near depth and the other at a far depth, suffice for distinguishing the two geometries. These two dots move in the same direction in the R geometry, and move in opposite directions in R+T geometry (Figure 2A and B). Notably, the viewing geometry (R vs. R+T) is not given directly to the network; it must infer the viewing geometry (and eye velocity) from optic flow and compute the output variables accordingly.

**Figure 8:**
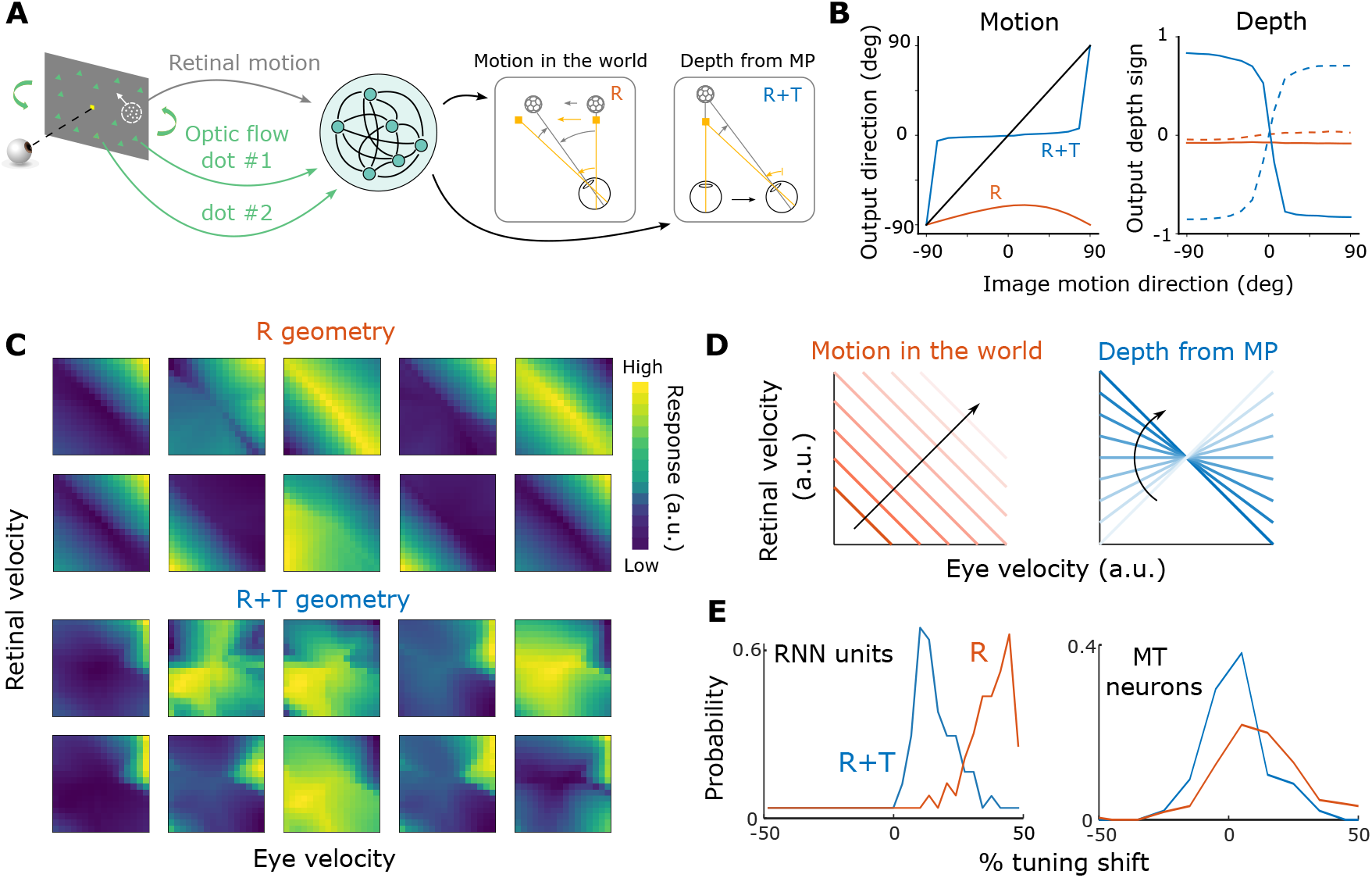
Recurrent neural network trained to perform motion and depth computations. **A**, Inputs and outputs of the network. The network receives three inputs—retinal motion of the target object and the image motion of two background dots (one near and one far relative to the fixation point)—and it produces two outputs, object motion in the world and depth from MP. **B**, Outputs of the trained RNN resemble human behavior. Left, The relationship between input retinal direction and the network’s estimated motion direction in R+T (blue) and R (orange) geometries. Right, Relationships between estimated depth sign and retinal direction. Dashed and solid curves: leftward and rightward eye movement, respectively. **C**, Joint velocity tuning for retinal and eye velocities in the R (top) and R+T (bottom) geometries for 10 example recurrent units. Corresponding units are shown for both geometries. **D**, Motion and depth computations require distinct joint representations of retinal and eye velocities. Left: velocity in world coordinates (orange lines) increases along the diagonal direction indicated by the black arrow. Right: depths from MP are represented by lines with varying slopes. **E**, Histograms showing distributions of tuning shifts observed in RNN units (left) and MT neurons (right; adapted from Xu and DeAngelis, 2022) for the R (orange) and R+T (blue) viewing geometries. A shift of 0% indicates a retinal-centered representation and a shift of 100% indicates a world-centered representation.

After training, the network’s behavior in both motion and depth perception tasks replicates the basic patterns observed in human data, showing a horizontal directional bias in the R condition, a vertical motion bias in the R+T condition, a nearly flat psychometric function for depth discrimination in the R condition, and robust depth discrimination performance that depends on eye direction in the R+T condition (Figure 8B, compare with Figures 6C–E and 7C–E). In addition, we found that recurrent units in the RNN show different joint tuning profiles for retinal velocity and eye velocity depending on the simulated viewing geometry. Specifically, a negative diagonal structure is more pronounced among recurrent units in the R geometry than in the R+T geometry (Figure 8C). This observation is consistent with the fact that the addition computation in the R context corresponds to a negative diagonal in the 2D joint velocity space (Figure 8D, left), whereas the division computation in the R+T geometry corresponds to lines with different slopes (Figure 8D, right). If neurons (or RNN units) are selective to a particular velocity in world coordinates, their joint velocity tuning would show a ridge along the negative diagonal (Figure 8C, top and Figure 8D, left); if neurons are selective to both a particular retinal motion and depth, their joint tuning would be a blob in the 2D space, as shown previously for some MT neurons (Figure 8C, bottom; Kim et al., 2017; Xu and DeAngelis, 2022).

We quantified the extent of the diagonal structure in these RNN units by examining the asymmetry in the 2D Fourier transform of the data (see Methods) and we compared the results with those found in MT neurons by Xu and DeAngelis (2022). Specifically, the percentage of tuning shift in the RNN units was measured as the normalized product of inertia in the 2D Fourier space (see Methods for details). We found that RNN units exhibit substantially more diagonal structure in the R geometry than R+T (∆median = 0.246, *p* = 7.755 × 10^*−*20^, Wilcoxon rank sum test; Figure 8E, left). Xu and DeAngelis (2022) modeled MT neurons’ responses to motion parallax stimuli under different conditions. While the experimental conditions were not the same as the R and R+T geometries discussed in this study, they have some similar features. When only retinal motion and eye movement were present, and no background optic flow was shown, the viewing geometry is likely to be more consistent with the R viewing geometry, in which a horizontal eye velocity is added to the retinal image motion. This is supported by similar patterns of results between the control and the R viewing geometry in the Pursuit condition of Experiment 1 (Figure 6D, gray dots versus orange dots). Xu and DeAngelis (2022) observed a significant shift in the tuning of MT neurons toward world coordinates during pursuit (median = 0.122, *p* = 2.44 × 10^*−*15^, one-tailed Wilcoxon signed-rank test; Figure 8E, right, orange). In contrast, when the animal fixated at the center of the screen and optic flow simulated the R+T geometry, the extent of the diagonal shift was significantly reduced (Z = 5.360, *p* = 4.17 × 10^*−*8^, one-tailed Wilcoxon rank sum test; Figure 8E, right, blue). These observations broadly align with our findings in RNN units, suggesting a potential role of MT neurons in flexibly computing object motion and depth under different viewing geometries.

## 4 Discussion

We demonstrate that the traditional model of visual compensation for pursuit eye movements, based on vector subtraction, fails to generalize to even the simplest combinations of eye translation and rotation. Instead, we provide a theoretical framework that relates object motion and depth (in the world) to retinal and eye velocities of a moving observer, across a range of possible viewing geometries. This framework unifies two well-known perceptual phenomena—coordinate transformation and computation of depth from motion parallax—that have generally been studied separately. We generated theoretical predictions for how perception of object motion and depth should depend on viewing geometry simulated by optic flow, and we verified these predictions using a series of well-controlled psychophysical experiments. Our results suggest that humans automatically, without any feedback or training, infer their viewing geometry from visual information (i.e., optic flow) and use this information in a context-specific fashion to compute the motion and depth of objects in the world. They flexibly attribute specific components of image motion to either eye rotation or depth structure depending on the inferred viewing geometry. A recurrent neural network trained to perform the same tasks shows underlying representations that are somewhat similar to neurons in area MT, suggesting a potential neural implementation of the flexible computations of motion and depth.

In the traditional view of visual perception during eye movements, sensory consequences of self-generated actions are considered detrimental to perception and, therefore, should be suppressed (e.g., Matin, 1974). By contrast, our study demon-strates that humans utilize the visual consequences of smooth pursuit eye movements (i.e., optic flow) to infer their viewing geometry and adaptively compute the depth and motion of objects. It is precisely the sensory consequences of self-motion that provide rich information about the relationship between the observer and the dynamic 3D environment, allowing more accurate perception of the 3D world during active exploration (Gibson, 1977; Aloimonos et al., 1988; Warren, 2021).

### 4.1 Interactions between object motion and depth

In our experiments, we presented the object only to one eye and kept its size constant, such that the object’s depth was ambiguous and subject to influence from motion information. Presumably, each subject’s prior distribution over possible object depths might affect their depth perception differently (Ooi et al., 2006; Knill, 2007; Burge et al., 2010). This might account for some of the cross-subject variability observed in the depth discrimination task. To directly test the interactions between motion and depth, a future direction would be to include additional depth cues for the object and to examine how this affects perceived motion direction. When different levels of depth cues (e.g., congruent vs. incongruent with the MP cue) are introduced, we might expect different amounts of retinal motion to be explained away as depth from MP. For example, if binocular disparity cues always indicated that the object was in the plane of the visual display, we would expect a smaller vertical bias in perceived direction in the R+T geometry, as one should not attribute a horizontal component of image motion to depth.

In addition, Equation (2) shows that object motion and depth are underdetermined in the absence of other depth cues, even when the viewing geometry is unambiguous. In the R+T geometry, the motion-pursuit law applies only when the object is stationary (Equation 4). When *p*^*′*^ = 1 and *ω*_*obj*_ ≠ 0, object motion must be subtracted from retinal image motion in order to accurately compute depth from MP:

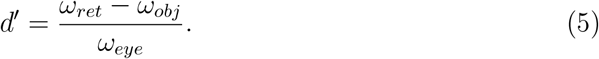

If the observer mistakenly believes the object is stationary and does not perform this subtraction, a bias in depth perception occurs. Indeed, this effect of object motion on depth perception was demonstrated in a recent study (French and DeAngelis, 2022).

A related line of research has investigated the phenomenon of optic flow parsing, namely the process of inferring object motion from optic flow (Warren and Rushton, 2007, 2008, 2009; MacNeilage et al., 2012; Foulkes et al., 2013; Niehorster and Li, 2017; Peltier et al., 2020). Warren and Rushton (2007) found that humans can differentiate between optic flow patterns generated by eye rotation (i.e., the R geometry) and lateral translation, showing that depth modulates object motion perception only in the latter case, as expected from the 3D geometry. Our study demonstrates that humans extract information about depth structure only when optic flow is depth-dependent, revealing a flexible and automatic switch between attributing the source of retinal motion to object motion vs. depth.

Previous studies have demonstrated the role of dynamic perspective cues (which are associated with rotation of the scene around the fixation point in the R+T geometry) in stabilizing the scene and disambiguating depth sign (Rogers and Rogers, 1992; Rogers, 2016; Buckthought et al., 2017), and neural correlates of depth coding from these visual cues have been found in macaque area MT (Kim et al., 2015). Our Experiment 2 provides the first direct evidence that humans automatically perceive depth from dynamic perspective cues in the R+T viewing geometry, with or without corresponding pursuit eye movements. Our findings thus add new insights into the visual processing of depth from MP.

### 4.2 Compensation for pursuit eye movement depends on viewing geometry

The perceptual consequences of pursuit eye movement have been extensively studied in the past decades (e.g., Filehne, 1922; Mack and Herman, 1973; Festinger et al., 1976; Wertheim, 1981, 1987; Swanston et al., 1992; Freeman and Banks, 1998; Freeman, 1999; Turano and Massof, 2001; Souman et al., 2005, 2006a,b; Spering and Gegenfurtner, 2007; Morvan and Wexler, 2009; Freeman et al., 2010; Spering and Montagnini, 2011; Furman and Gur, 2012). These studies often hypothesize that the brain generates a reference signal related to eye velocity and subtracts it from the retinal image motion in order to perceive a stable world (e.g., Freeman and Banks, 1998; Wertheim, 1987) (Figure 1C). While this line of research has successfully explained many visual phenomena that involve pure eye rotations in 2D displays, it does not generalize to situations in which there is 3D scene structure and combinations of eye translation and rotation (Figure 2). Eye translation introduces components of optic flow that are depth-dependent, such that one cannot perform a simple vector subtraction to compensate for the visual consequences of smooth pursuit. Our study provides a much more general description of how the brain should compute scene-relative object motion and depth under a variety of viewing geometries that involve combinations of eye translation and rotation.

Importantly, we observed similar patterns of perceptual biases in the Fixation and Pursuit conditions, suggesting that the different effects of viewing geometry on motion and depth perception cannot simply be explained by retinal slip caused by imperfect pursuit eye movements. Interestingly, in the Pursuit conditions, the effects are similar between the R condition and the no background condition (Figure S1, orange vs. gray markers), indicating that either real pursuit alone or optic flow alone can shift perceived object direction toward the eye movement. This suggests an absence of additive effects between extraretinal signals and optic flow in the CT computation. However, our experiments were not designed specifically to quantify the relative contributions of optic flow and extra-retinal signals to inferring viewing geometry, and this would be a valuable topic for further research.

A few previous studies have investigated the interactions between motion and depth perception during self-motion. For example, Wallach et al. (1972) showed that underestimating an object’s depth resulted in an illusory rotation of the object during lateral head translation. Gogel and colleagues investigated how this apparent motion changed as a function of under- or over-estimation of depth, showing that the direction of the perceived motion changed systematically based on the geometry of motion parallax (Gogel and Tietz, 1973; Gogel, 1980). In addition, illusory motion induced by head motion can be added to or subtracted from the physical movement of objects (Gogel, 1979; Gogel and Tietz, 1979; Rogers, 2016).

In these studies, observers were explicitly instructed to laterally translate their heads (Gogel, 1976). As a result, the viewing geometry was always unambiguous and the role of different geometries was not explored. The source of uncertainty, or errors, was thought to be either the intrinsic underestimation of distance in a dark room (Wallach et al., 1972; Gilinsky, 1951; Gogel, 1969) or that induced by binocular disparity cues (Gogel, 1980). How does the observer’s belief about the viewing geometry modulate the interaction between motion and depth? What are the cues (e.g., optic flow and extraretinal signals) for disambiguating viewing geometry? These important questions have not been addressed by previous studies.

We demonstrate for the first time that humans use optic flow information to infer their viewing geometry and flexibly compute object motion and depth based on their interpretations of the geometry. Moreover, our work provides novel insights into how the brain solves the causal inference problem of parsing retinal image motion into different causes—object motion in the world and depth from motion parallax—based on the information about self-motion given by optic flow.

### 4.3 Recurrent neural network and the neural basis of contextual computation

Our RNN model provides insights into the neural basis of computing object motion and depth in different viewing geometries. By comparing representations in the network model with neurons in MT, we suggest a potential role of MT neurons in implementing flexible computations of motion and depth. Area MT has been linked to the perception of object motion (e.g., Britten et al., 1992, 1996; Albright, 1984), and perception of depth based on both binocular disparity (e.g., DeAngelis and Newsome, 1999; DeAngelis et al., 1998) and MP cues (e.g., Nadler et al., 2008; Kim et al., 2015). Emerging evidence shows that sensory areas receive top-down modulations from higher cortical regions that reflect perceptual decision variables or cognitive states (e.g., Bondy et al., 2018; Keller et al., 2020). Therefore, we speculate that neurons in MT might receive feedback signals about viewing geometry from higher-level areas, such as the medial superior temporal (MST) area or areas in the intraparietal sulcus, and use these signals to modulate the response to retinal motion. A recent study has shown that neurons in dorsal MST are selective for large-field optic flow that simulates eye translation and rotation in the R+T geometry (DiRisio et al., 2023), which suggests a potential source of information about viewing geometry that is known to feed back to area MT (Maunsell and van Essen, 1983; Ungerleider and Desimone, 1986; Felleman and Van Essen, 1991). Furthermore, a recent study Peltier et al. (2024) has demonstrated that responses of MT neurons are modulated by background optic flow in a manner that is consistent with perceptual biases that are associated with optic flow parsing (i.e. flow parsing; Warren and Rushton, 2007, 2008, 2009; MacNeilage et al., 2012; Foulkes et al., 2013; Niehorster and Li, 2017; Peltier et al., 2020). However, whether activity in area MT will reflect flexible computations of object motion and depth, based on inferred viewing geometry, remains to be examined. Another possible source of signals related to viewing geometry, as suggested by our RNN model, is the recurrent connections within area MT. Different optic flow patterns used in our study might differentially trigger responses of a subset of MT neurons whose receptive fields overlapped with the background dots, and these responses could, in turn, modulate MT neurons with receptive fields overlapping the object. As shown by Xu and DeAngelis (2022), a partial shift in the tuning preference observed in MT neurons, in theory, suffices for computing world-centered motion and depth. Further investigation with inactivation techniques would be desirable to determine whether or not higher-level cortical areas are involved in these flexible computations of motion and depth.

### 4.4 Limitations and future directions

Deriving from 3D geometric principles, our theoretical framework makes quantitative predictions about motion and depth perception during self-motion for a range of scenarios as depicted in Figure 3. Perceptual biases, as predicted by our framework, were demonstrated for perception of object motion and depth in two simple viewing geometries: Pure Rotation (R) and Rotation + Translation (R+T). In addition, our framework also makes predictions for scenarios in which the viewing geometry is intermediate between R and R+T. Examination of these intermediate viewing geometries in future studies will provide further validation of our theory.

A limitation of our theory is that Equation (2) only applies when both the observer and the pursuit target translate in the fronto-parallel plane, such that the relative position of the rotation pivot remains constant (Figure 3). Because natural behavior involves movements along multiple axes at the same time (Matthis et al., 2022), an extension of our theory is needed to better understand visual perception in natural environments. Further analysis of how optic flow is constrained by different viewing geometries might yield insights into a more generalized theory (Longuet-Higgins and Prazdny, 1980; Thompson and Pong, 1990; Nelson, 1991).

Although our psychophysical data broadly align with our theoretical predictions, they are not fully accounted for by Equation (2). Specifically, the measured perceptual biases are typically partial biases. Here, we capture substantial deviations from the ideal predictions using discount factors analogous to a flow parsing gain (Niehorster and Li, 2017). However, multiple sources might contribute to these deviations: observation noise, uncertainty about the viewing geometry, underestimation of smooth pursuit eye velocity (Festinger et al., 1976), cue conflict between vestibular and visual signals (Dokka et al., 2015), a slow-speed prior (Stocker and Simoncelli, 2006), and so on. To fully quantify and understand the inference performed by the observer, a more comprehensive probabilistic model of motion and depth perception will be needed (Gershman et al., 2016; Shivkumar et al., 2023). Specifically, the problem of differentiating between object motion, depth, and self-motion might be formalized as an instance of the Bayesian causal inference problem (Kording et al., 2007; Shams and Beierholm, 2010; Dokka et al., 2019; French and DeAngelis, 2020).

In our neural network simulations, the RNN was directly trained to reproduce the predicted motion and depth perception. Whether or not such perceptual biases naturally emerge in networks trained to estimate self-motion or encode videos of natural scenes is an interesting future direction to explore (Mineault et al., 2021; Vafaii et al., 2024). Our comparison of MT neural responses between the R and R+T conditions was indirect, as previous experimental work did not explicitly simulate these two viewing geometries (Nadler et al., 2008, 2009; Kim et al., 2015; Xu and DeAngelis, 2022). An ongoing study that directly measures the responses of MT neurons in the two viewing geometries will provide new insights into the neural mechanisms underlying flexible computations of motion and depth.

## 5 Methods

### 5.1 Participants

Ten participants (4 males and 6 females, 18–58 years old) with normal or corrected-to-normal vision were recruited for the psychophysical experiments. All participants had normal stereo vision (<50 arcseconds, Randot Stereotest). Seven of the participants were naive to the experiments and unaware of the purpose of this study. All participants completed the depth discrimination task, and nine of them finished the motion estimation task. Informed written consent was obtained from each participant prior to data collection. The study was approved by the Institutional Review Board at the University of Rochester.

### 5.2 Apparatus

Participants sat in front of a 48-inch computer monitor (AORUS FO48U; width, 105.2 cm; height, 59.2 cm) at a viewing distance of 57 cm, yielding a field of view of ∼ 85^°^×55^°^. A chin and forehead rest was used to restrict participants’ head movements. Position of each participant’s dominant eye was monitored by an infrared eye tracker (Eyelink 1000Plus, SR-Research) positioned on a desk in front of the subject at a distance of ∼52 cm. The refresh rate of the monitor was 60 Hz, the pixel resolution was 1920×1080, and the pixel size was ∼ 2.6^*′*^ × 3^*′*^ arcmin. During the experiments, participants viewed the visual display through a pair of red-blue 3D glasses in a dark room. The mean luminance of the blank screen was 0 cd/m^2^ (due to the OLED display), the mean luminance of the object was 1.383 cd/m^2^, and the mean luminance of the background optic flow was 0.510 cd/m^2^.

### 5.3 Stimuli

Visual stimuli were generated by custom software and rendered in 3D virtual environments using the OpenGL library in C++. Participants were instructed to fixate on a square at the center of the screen, and a random-dot patch (referred to as the “object”; radius 8^°^) was presented on the horizontal meridian, interleaved between left and right hemifields, at 10^°^ eccentricity. Viewing of the object was monocular to remove binocular depth cues. The size of the dots comprising the object was constant on the screen across conditions, to avoid providing depth cues from varying image sizes. In most conditions, a full-field 3D cloud of background dots was presented for the same duration as the object. The motion of the background dots was generated by moving the OpenGL camera, simulating either the R or R+T viewing geometries (Figure 2A and B; see Supplementary Information for details). The movements of the object, background optic flow, and fixation target followed a Gaussian velocity profile (±3*σ*) spanning a duration of 1 s. The immediate region surrounding the object (2× the object’s radius) was masked to avoid local motion interactions between background dots and the object.

#### Simulating viewing geometry with optic flow

A 3D cloud of background random dots simulated 4 different configurations of eye translation and/or rotation. 1) In the R Fixation condition (Video 3), the OpenGL camera rotated about the y-axis (yaw rotation) to track the moving fixation target such that it remained at the center of the screen; this resulted in rotational optic flow that simulated the R geometry, while requiring no actual pursuit eye movement. 2) In the R+T Fixation condition (Video 4), the OpenGL camera translated laterally while counter-rotating to keep the world-fixed fixation target at the center of the screen. This generated background optic flow that simulated both translation and rotation in the R+T geometry, while again requiring no smooth pursuit. 3) In the R Pursuit condition (Video 5), the OpenGL camera remained stationary. As a result, the background dots did not move on the screen. A fixation target appeared at the center of the screen and moved 3 cm, either leftward or rightward. The movement of the fixation target followed a Gaussian speed profile (±3*σ*) spanning 1 second, resulting in a peak speed of ∼13^°^/s and a mean speed of ∼ 5.3^°^/s. Subjects were required to track the fixation target with their eyes. 4) In the R+T Pursuit condition (Video 6), the OpenGL camera translated laterally (leftward or rightward) by 3 cm in the virtual environment (following the same Gaussian speed profile). Therefore, the background dots appeared to translate in the opposite direction on the screen, providing optic flow that simulated eye translation. Throughout the trial, a fixation target appeared at a fixed location in the virtual environment but moved on the screen due to the camera’s translation. The subject was required to make smooth eye movements to remain fixated on the world-fixed target. Note that background elements were triangles of a fixed size in the scene, such that their image size was inversely proportional to their distance (unlike for the object stimulus).

#### Object motion

A random-dot patch (the “object”) with a fixed dot density of ∼ 4.7 dots/^°^ and a dot size of ∼ 0.2^°^ was rendered monocularly to the right eye. To make the object’s depth ambiguous, the dot size and diameter of the aperture were kept constant across all stimulus conditions. In each trial, the object moved as a whole (the aperture and the random dots within it moved together), in one of several directions on the screen. The position and motion trajectory of the object were carefully computed such that it yielded identical image motion between the R and R+T conditions (see Supplementary Information for details). The speed of the object followed a Gaussian profile with a maximum speed of ∼ 6.67^°^/s and a mean speed of ∼ 2.67^°^/s.

### 5.4 Experiment 1: Procedures and experimental conditions

In Experiment 1, subjects performed a motion estimation task. At the beginning of each trial, a fixation point appeared at the center of the screen, followed by the onset of the object and background dots. In each trial, the direction of retinal motion of the object was randomly chosen from 0^°^ to 360^°^ with 30^°^ spacing (we defined the rightward direction as 0^°^ and the angle increases in a counter-clockwise direction). The object and background were presented for 1 second, after which another patch of dots (the “probe”; rendered binocularly at the same depth as the screen) appeared at the same location on an otherwise blank screen. Participants used a dial to adjust the motion direction of the probe such that it matched the perceived direction of the object. After adjusting the dial, participants pressed a button to register their response and proceeded to the next trial after an inter-trial interval of 1.5 seconds. Failure to register the response within 5 seconds from probe onset resulted in a time-out, and the trial was repeated at a later time. Eye position was monitored throughout each trial, and failure to maintain fixation within a ±5^°^ rectangular window around the fixation target resulted in a failed trial, after which visual stimuli would be immediately turned off. Audio feedback was provided at the end of each trial to indicate successful completion of the trial with a high-pitched tone and failed trials (fixation break or time out) with a low-pitched tone. For completed trials, information about response error was not provided to the participants in any form. This lack of feedback prevented participants from learning to compensate for perceptual biases induced by optic flow.

Four main stimulus conditions were presented: two eye-movement conditions × two background conditions (Figure 4; Videos 3–6). The two eye-movement conditions were: 1) the Pursuit condition, in which the subject visually tracked a fixation target that moved across the center of the screen while simultaneously viewing an object composed of random dots at 10^°^ eccentricity; 2) the Fixation condition, in which participants fixated on a stationary target at the center of the screen while background dots simulated eye translation and/or rotation. The direction of actual or simulated eye movements was either 0^°^ (rightward) or 180^°^ (leftward), randomly interleaved across trials. The two background conditions were: 1) the R viewing geometry, in which the motion of the background dots was consistent with a pure eye rotation; 2) the R+T viewing geometry, in which the background dots simulated a combination of lateral translation and rotation of the eye. In addition, two control conditions were interleaved with the main conditions, including Pursuit and Fixation conditions with object motion in the absence of background dots (Videos 7 and 8).

Before the main experimental session, a practice session (72–144 trials) was completed to ensure a correct understanding of the task and to give subjects practice with the dial-turning behavior. In this short block of practice trials, only the object was present and no background was shown. All subjects successfully reported the object’s motion direction within a ±15^°^ range around the ground truth before proceeding to the main experimental session.

### 5.5 Experiment 2: Procedures and experimental conditions

In Experiment 2, subjects performed a depth discrimination task. The visual stimuli and experimental procedure were the same as in Experiment 1, except that participants pressed one of two buttons, either during the stimulus period or within 2.5 seconds afterward, to report whether the object was located near or far compared to the fixation point. Background and eye movement conditions were the same as those in Experiment 1. In each trial, the direction of retinal motion of the object was randomly chosen from 0^°^ to 180^°^ with 15^°^ spacing. Because depth is expected to be determined by the horizontal component of retinal motion and eye velocity, we did not include directions in the range between 180^°^ to 360^°^, which differ from 0^°^–180^°^ only in vertical components.

Before the formal experimental session, participants underwent a practice session to become familiar with the stimuli and the task. In the practice session, the background motion was the same as the R+T condition, and the object moved in horizontal directions at different speeds such that its retinal motion could be fully explained as depth from motion parallax. After 72 practice trials, the experimenter decided to either 1) proceed to the formal experimental session if the accuracy was above 95%, or 2) continue with another practice session in which the object was viewed binocularly to aid depth perception. After the binocular session, another monocular practice session was run to ensure that participants performed the task well above chance. Three subjects did not proceed to the formal experimental sessions due to failure to report depth at an accuracy above 80% during the practice sessions.

### 5.6 Data analysis

#### Analysis of eye-tracking data

For most subjects, eye position signals measured by the eye-tracker were used to ensure fixation behavior and to compute smooth pursuit gains. In the fixation conditions, trials with eye positions outside of a 10^°^-by-10^°^ rectangular window around the fixation target for over 100 ms were excluded from the analysis. In the pursuit conditions, pursuit velocities were obtained by filtering the eye position data with a first-derivative-of-Gaussian window (SD = 25 ms), followed by a velocity threshold at 40^°^/s and an acceleration threshold at 300^°^*/s*^2^ to remove catch-up saccades and artifacts. The median of the pursuit velocity was obtained across trials, and the ratio between its peak and the peak velocity of the pursuit target was computed as pursuit gain. Across all subjects, the mean pursuit gains are 0.755 (SE = 0.119) in Experiment 1 and 0.768 (SE = 0.0512) in Experiment 2 (Table 1). There was no significant difference between the pursuit gains in Experiments 1 and 2 (*p* = 0.535, Wilcoxon rank sum test). Across subjects, pursuit gain was not correlated with *a*_*ret*_ and *a*_*eye*_ in Experiment 1 (*r* = 0.586, *p* = 0.097 for *a*_*ret*_ and *r* = −0.291, *p* = 0.447 for *a*_*eye*_ in the R geometry; *r* = −0.392, *p* = 0.296 for *a*_*ret*_ and *r* = 0.104, *p* = 0.789 for *a*_*eye*_ in the R+T geometry) or the magnitudes of slopes in Experiment 2 (*r* = −0.092, *p* = 0.844 for the R geometry and *r* = −0.528, *p* = 0.230 for the R+T geometry).

**Table 1.** Pursuit gains of each participant in Experiments 1 and 2. Eye tracking was not conducted for h201 and h500 in Experiment 1, and the Pursuit conditions were not included in Experiment 2 for h201, h500, and h501. Participant h508 did not participate in Experiment 1.

| Participant | Experiment 1 | Experiment 2 |
| --- | --- | --- |
| h201 | / | / |
| h500 | / | / |
| h501 | 1.376 | / |
| h507 | 0.436 | 0.569 |
| h508 | / | 0.859 |
| h510 | 0.652 | 0.622 |
| h512 | 0.736 | 0.758 |
| h518 | 0.731 | 0.816 |
| h520 | 0.876 | 0.963 |
| h521 | 0.476 | 0.787 |

#### Behavioral data analysis

In Experiment 1, because we expect the pattern of biases to be symmetric around the horizontal axis (Figure 2), image motion directions and reported directions from trials in which the object moved in directions from 180^°^–360^°^ were flipped horizontally and pooled with those from trials with retinal directions in the range of 0^°^–180^°^. Following a similar logic, we pooled data from trials with rightward and leftward eye movements by flipping the velocities vertically. This results in a consistent leftward bias prediction in the R geometry shown in Figure 6C.

Because of imperfect pursuit eye movements by the participants, the actual retinal image motion of the object was contaminated by retinal slip. It differed from the intended velocity in the Pursuit conditions. We corrected this by factoring in the measured pursuit gain, *g*_*pursuit*_, for each subject:

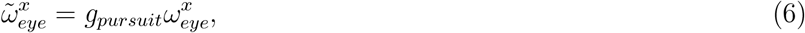

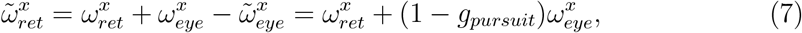

where 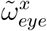 and 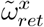 are the horizontal components of the real eye and retinal velocities, respectively; 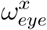 and 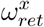 are the intended horizontal components of eye and retinal velocities, respectively. Because pursuit eye movements were always along the horizontal axis, the vertical components of the velocities were unaffected.

In the R+T viewing geometry, we assumed that a portion of the horizontal component of retinal image velocity would be explained as motion parallax for computing depth:

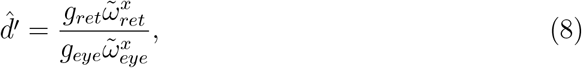

where 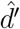 is the perceived relative depth and *g*_*ret*_ represents the proportion of horizontal retinal motion perceived as motion parallax. Similarly, we assumed that a portion of eye velocity was accounted for by a factor, *g*_*eye*_, in the R viewing geometry. Therefore, Equation (2) can be rewritten as:

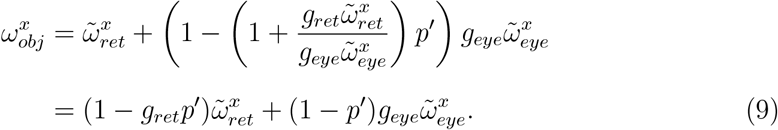

This formula indicates that perceived object motion is a linear combination of retinal and eye velocities, with varying weights on each velocity term that depend on the viewing geometry, *p*^*′*^. Simplifying this equation, we used a linear model to capture this relationship:

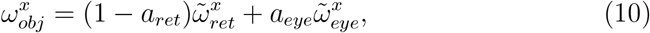

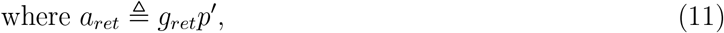

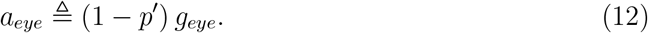

In the R geometry, *p*^*′*^ = 0, and we expect *a*_*ret*_ = 0, *a*_*eye*_ *>* 0; by contrast, in the R+T geometry, *p*^*′*^ = 1, and we expect *a*_*ret*_ *>* 0, *a*_*eye*_ = 0. To test this prediction, this linear model was fit to the direction reports in each of the conditions by minimizing the mean cosine error with L1 regularization to impose sparsity:

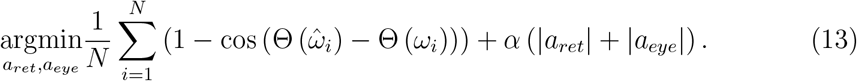

Here, 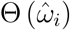 and Θ (*ω*_*i*_) indicate the predicted and actual reported object motion directions in the *i*-th trial. Regularization strength, *α*, was chosen by cross-validation. Optimization was done using the *fminsearch* function in MATLAB (Mathworks, MA). *a*_*ret*_ and *a*_*eye*_ were bounded in the range of [0,1].

For Experiment 2, a cumulative-Gaussian psychometric function was fit to binary depth reports in each viewing geometry using the *psignifit* library (Schutt et al., 2016) in MATLAB:

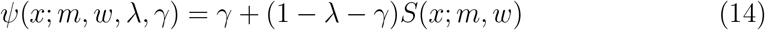

where *λ* and *γ* denote the lapse rate at the highest and lowest stimulus levels. *S* is the cumulative Gaussian function:

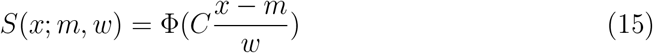

where *x* is the retinal direction of the object, *m* and *w* are the mean and standard deviation of the Gaussian function, respectively, and *C* = Φ^*−*1^(0.95) − Φ^*−*1^(0.05). Confidence intervals around parameters were obtained by bootstrapping inference provided in the *psignifit* library (Schutt et al., 2016).

### 5.7 Recurrent neural networks and neural data

#### Architecture

RNN models were implemented using the PsychRNN library (Ehrlich et al., 2021) and Tensorflow (Abadi et al., 2015). The RNN consisted of three input units, 64 recurrent units, and two output units. Our goal is to model the inputs and outputs relevant to the psychophysical experiment while keeping the network structure simple. For inputs, we use one input unit to represent the horizontal component of the object’s retinal motion, and the other two units to represent the horizontal components of background optic flow for two different depths. We reasoned that the minimum information needed to disambiguate the viewing geometry (R vs. R+T) is the flow vector of two background dots, one at a near depth and the other at a far depth. The two outputs of the network were scalars representing the horizontal component of the object’s motion in the world and its depth (positive means far and negative means near). Recurrent units were fully connected; each unit received all inputs and was connected to both outputs. The dynamics of the network can be described as:

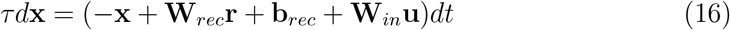

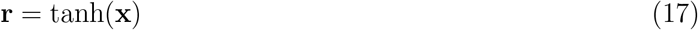

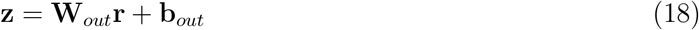

where **u, r, x**, and **z** denote the input, activation of the hidden layer, recurrent state, and output. *τ* and *dt* are the predefined time constant and time step, respectively, and *τ* = 100 ms, *dt* = 10 ms . **W**_*in*_, **W**_*rec*_, and **W**_*out*_ are the learnable weight matrices for input, recurrent, and output connections. **b**_*rec*_ and **b**_*out*_ are biases fed into the recurrent and output units.

#### Task and training

In the initial 500-ms period of each simulated trial, the values of the inputs were independent Gaussian noise, 𝒩 (0, 0.5), representing sensory noise. After that, the stimulus was presented for 1 s, represented by a constant value of retinal motion, optic flow vector of a near dot, and flow vector of a far dot, in addition to the Gaussian noise. The scale of Gaussian noise was chosen to qualitatively match the slopes of the model’s psychometric curves (Figure 8B) to those observed in human participants (Figure S2). The stimulus presentation period was followed by another 500 ms of noise. The output channels corresponded to the horizontal velocity of the object in world coordinates and depth from MP, and the network was trained to minimize the total L2 loss on these outputs only during the last 500 ms of each trial, after stimulus presentation was completed. Optimization was done with the ADAM optimizer (Kingma and Ba, 2014) implemented in TensorFlow (Abadi et al., 2015). There were 50,000 training epochs, and the learning rate was 1 × 10^*−*3^. The batch size was 128. In each trial, the retinal and eye velocities were uniformly sampled from a range of -10 to 10 (arbitrary units).

#### Psychometric functions of the RNN

After training, psychometric functions for the motion estimation and depth discrimination tasks were obtained by running predictions of the RNN on a set of inputs that replicated the human psychophysical experiments. Retinal motion directions ranged from 0^°^to 180^°^with a spacing of 12^°^and the speed was constant at 2 (arbitrary units). The speed of eye velocity was 3 times that of the retinal motion and the directions were leftward and rightward. Horizontal components of the retinal motion and eye velocity were used as inputs, and the model’s estimated object motion direction was obtained by taking the arctangent between the veridical vertical component of the object’s motion and the model’s estimate of its horizontal speed.

#### Tuning of single units in the network

After training, we tested the RNN on a grid of stimuli covering all retinal and eye velocity combinations ranging from -10 to 10 with a spacing of 1 (arbitrary units) and both viewing geometries. For each recurrent unit, the joint velocity tuning profile at each time point was obtained by mapping the activation of the test stimuli to the 2D velocity grid.

#### Tuning shifts in recurrent units

We quantified the extent of tuning shifts in each joint tuning profile as the degree of asymmetry in its 2D Fourier transform. A shift of retinal velocity tuning with eye velocity would manifest as a diagonal structure in the joint tuning profile (Xu and DeAngelis, 2022), and such diagonal structures will produce an asymmetric 2D Fourier power spectrum (DeAngelis et al., 1993). Specifically, we took the 2D Fourier transform of the joint tuning profile at the last time point for each recurrent unit of the network and thresholded the power spectrum at -10 dB to reduce noise. We then computed the normalized product of inertia of the power spectrum as

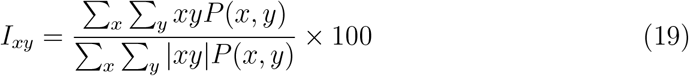

where *x* and *y* are coordinates in the 2D Fourier domain, and *P* (*x, y*) is the power at (*x, y*). The normalized product of inertia ranges from -100% to 100%, with 0% indicating no tuning shift, 100% being maximally shifted towards world coordinates, and -100% showing maximum tuning shifts in the opposite direction of world coordinates. This metric allows us to quantify the extent of tuning shifts without assuming a specific form of the joint tuning profile; therefore, it is more generally applicable than our previous measure using parametric model fitting (Xu and DeAngelis, 2022).

#### Tuning shifts in MT neurons

Due to the limited samples in the 2D velocity space of the experimental data in Xu and DeAngelis (2022), we could not use the normalized product of inertia to measure tuning shifts in the neural responses of MT neurons. Instead, we used the estimated weights on eye velocity developed to measure the tuning shifts in MT neurons (Xu and DeAngelis, 2022). In brief, we modeled neural responses to eye velocity and retinal motion as a combination of tuning shift, multiplicative gain, and additive modulation:

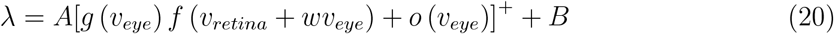

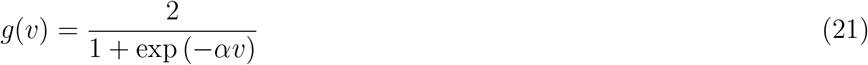

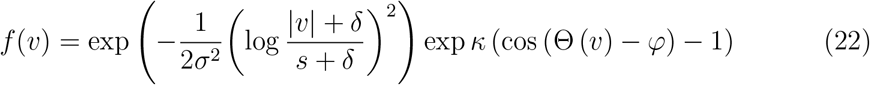

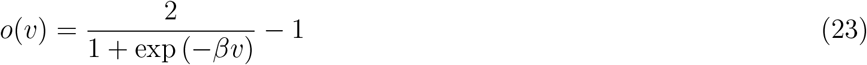

Here, *λ* is the estimated firing rate; *v*_*retina*_ and *v*_*eye*_ are retinal and eye velocities at each time point; *A* and *B* are the amplitude and baseline firing rate; *w* is the weight on eye velocity that quantifies the extent of tuning shifts; [·]^+^ is a rectifier that prevents negative firing rates; *g*(*v*) is the multiplicative gain function; *α* controls the slope of the gain function; *f* (*v*) is the tuning function; *σ, δ, s, κ*, and *φ* jointly define the width and offset of the function; |*v*| and Θ(*v*) denote the speed and direction of the velocity; *o*(*v*) is the additive modulation function, and *β* controls the slope of the additive function. Free parameters in the model were estimated by minimizing the negative log-likelihood assuming Poisson noise.

The estimated weights on eye velocity, *ω*, range from -1 to 1, with 0 being retinal-centered, 1 being completely world-centered, and -1 being the opposite of the expected shift. While strictly speaking, this measure is not equivalent to the normalized product of inertia used for hidden units, they are bounded in the same range and are roughly linearly related. Therefore, we used them as measures of tuning shifts and compared the distributions of these metrics between RNN units and neurons in MT.

## 8 Supplementary Information

**Figure S1:**
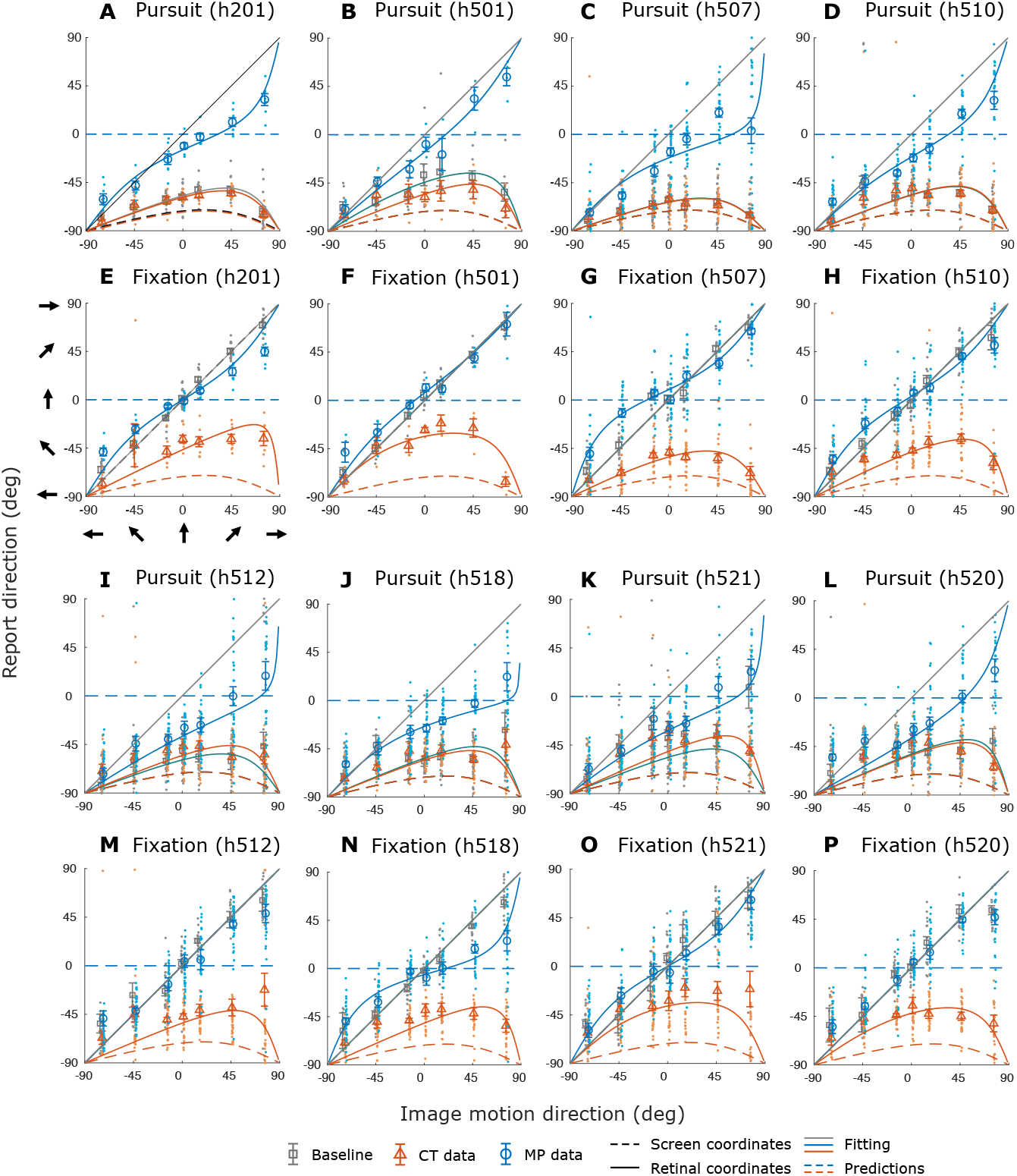
Results of the motion estimation task from additional participants. **A-D & I-L**, Data from the Pursuit condition for four additional participants. **E-H & M-P**, Data from the Fixation condition for the same participants. Format as in Figure 6D and E.

**Figure S2:**
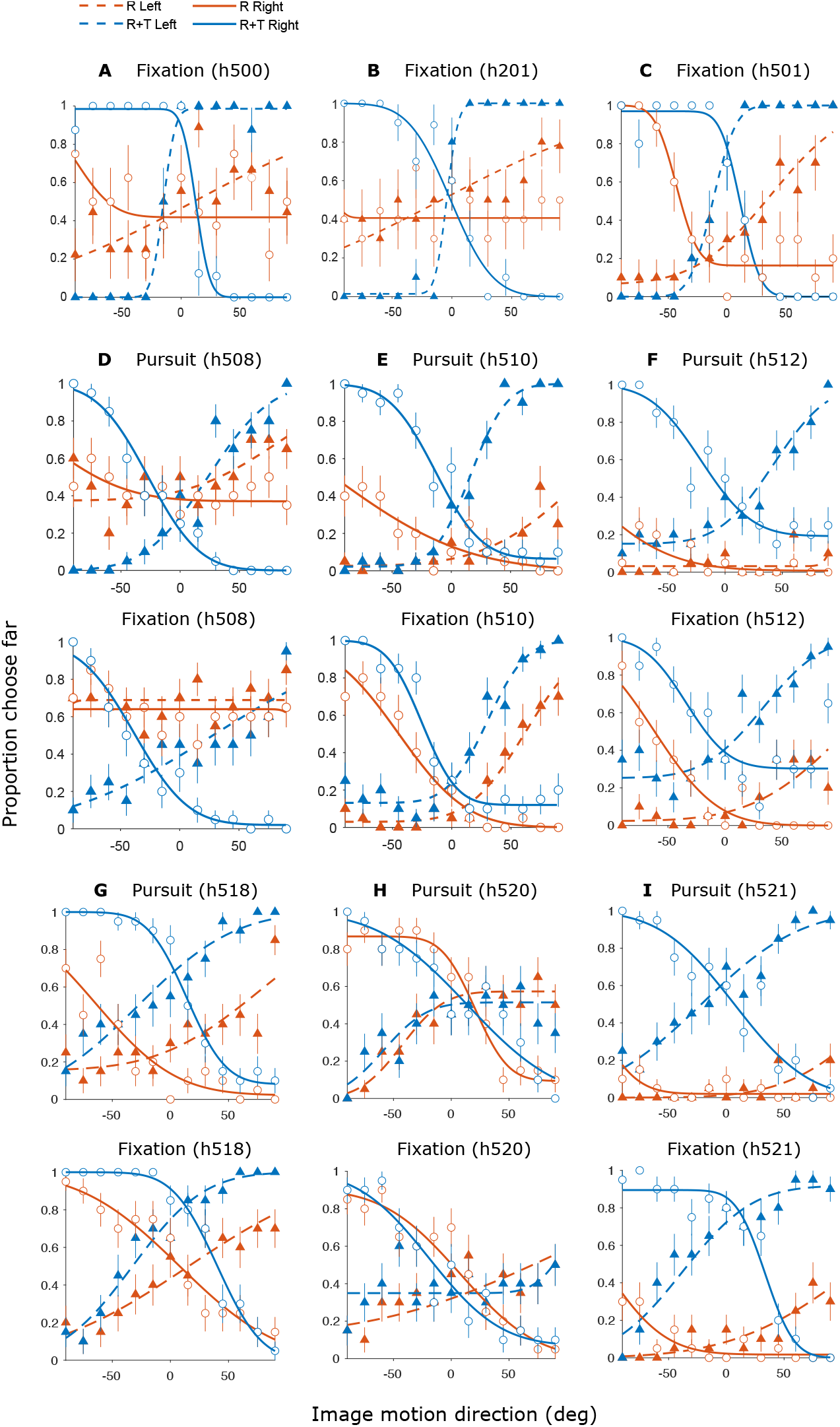
Results from the depth discrimination task for additional participants. **A-C**, Results from three non-naive participants in the Fixation condition. **D-I**, Results from naive participants in the Pursuit (top) and Fixation (bottom) conditions. Format as in Figure 7D and E.

### 8.1 Videos

Videos of the stimuli can be found at https://osf.io/zy8w6/?view_only=894cfd40b92e437586e29dc3d1be5441.

Video 1. Optic flow in the pure rotation (“R”) viewing geometry.

Video 2. Optic flow in the rotation + translation (“R+T”) viewing geometry.

Video 3. Stimulus for the R Fixation condition. Optic flow simulates rightward eye rotation and the object moves toward the top-right corner.

Video 4. Stimulus for the R+T Fixation condition. Optic flow simulates rightward eye rotation and leftward eye translation; the object moves in the same direction as in Video 3.

Video 5. Stimulus for the R Pursuit condition. The fixation target moves rightward and the object moves toward the top-right corner. The object’s motion direction relative to the fixation target is the same as in Videos 3 and 4.

Video 6. Stimulus for the R+T Pursuit condition. The fixation target moves rightward, optic flow simulates a leftward translation, and the object moves in the same direction as in Video 5.

Video 7. Stimulus for the control Fixation condition. No background dots were present and the object moves in the same direction on the screen as in Videos 3 and 4.

Video 8. Stimulus for the control Pursuit condition. No background dots were present and the object moves in the same direction on the screen as in Videos 5 and 6.

### 8.2 Supplementary figures

### 8.3 Derivation of viewing geometry

Consider a general scenario in which the observer translates their body (or head) laterally while tracking a moving fixation target by pursuit eye movements with an angular velocity relative to the scene, *ω*_*eye*_ (Figure 3). Meanwhile, another object moves independently in the frontoparallel plane at a certain distance, *d*, from the fixation target.

The retinal motion, *ω*_*ret*_, of the object has three components: (1) image motion produced by object motion in the world, *ω*_*obj*_, (2) motion parallax produced by observer’s translation, 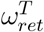, and (3) image motion produced by the eye rotation that tracks the moving fixation target, 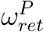 (Figure 3):

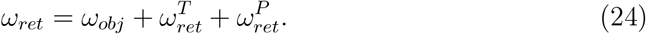

Rewriting the approximate form of the motion-pursuit law (Nawrot and Stroyan, 2009), 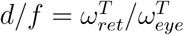, the motion parallax component is computed as:

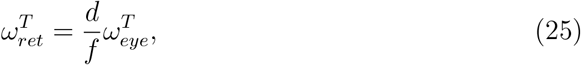

where *f* is the viewing distance and 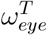 is the angular velocity of the eye rotation (relative to the scene) that compensates for the eye’s translation relative to the scene (as opposed to the eye rotation needed to track a moving fixation target).

To obtain 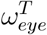, consider the intersection at a distance, *p*, between the line of sight at the initial time point *t*_0_ and that at a later time point *t*_0_ + *dt*. The position of *p* describes the relationship between the movement of the fixation target and that of the observer, and 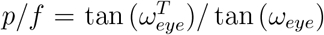, where *ω*_*eye*_ is the angular velocity of the total eye rotation. For small angles, tan(*ω*) ≈ *ω*, thus 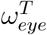 can be computed as:

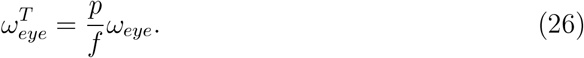

Note that while the distance between the rotation pivot and the eye, *p*, can change as the eye translates, the ratio between *p* and *f* remains constant for lateral translations (Figure 3).

The third component of retinal motion, 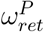, is the opposite of the eye velocity caused by a moving fixation target, 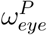 :

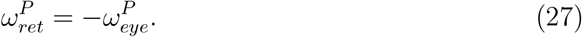

Because 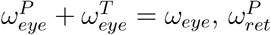 can be computed as:

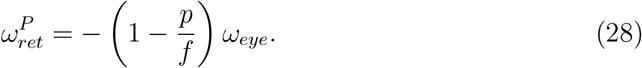

From Equations (24) to (28), we can obtain the angular velocity of the object as:

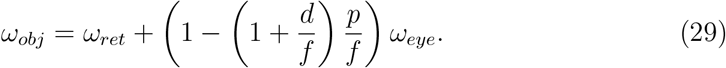

Normalizing the object’s depth, *d*, and the rotation pivot, *p*, by viewing distance, *f*, we have:

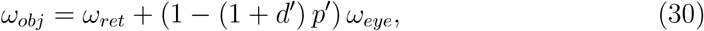

where *d*^*′*^ ≜ *d/f* and *p*^*′*^ ≜ *p/f* . In the absence of another depth cue, *ω*_*obj*_ and *d*^*′*^ are underdetermined, even if *p*^*′*^ is specified.

Notably, although we use the approximate formula for the motion-pursuit law here (Equation 4; Nawrot and Stroyan, 2009), Equation (2) still applies if we replace *d*^*′*^ with a more accurate form, *d*^*′*^ ≜ *d/*(*d* + *f*). It is also worth noting that we only consider scenarios in which the observer, pursuit target, and object translate in the fronto-parallel plane, as depicted in Figure 3. When the pursuit target moves in depth, the rotation pivot *p*^*′*^ is not constant and Equation (2) no longer applies.

### 8.4 Details of stimulus generation

To ensure that the motion of the object on the screen was the same across the two viewing geometries, we derived the relationship between the simulated 3D geometry in OpenGL and the screen projections based on standard projective geometry. The 3D coordinates of the object and camera were then determined by back-tracing from the desired image positions and motion.

#### Projective geometry

3D coordinates in a virtual OpenGL environment can be converted to a normalized screen coordinate system in two steps (Woo et al., 1999).

First, multiply the 3D coordinates, 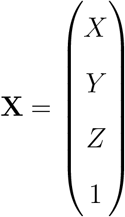, with a perspective projection matrix, **P**:

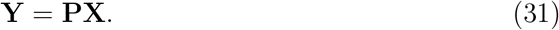

Here, the X-axis is pointing from left to right, Y-axis is pointing upward, Z-axis is pointing forward, and the projection matrix is given by:

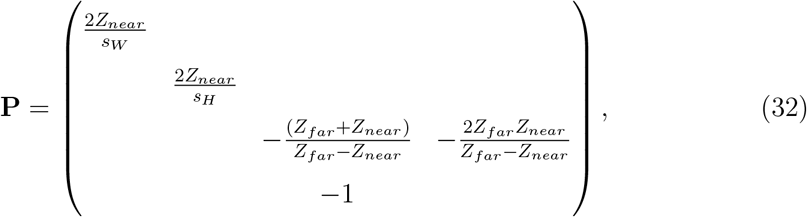

where *Z*_*near*_ and *Z*_*far*_ are the z-coordinates of the near and far clipping planes, respectively. In our experiments, *Z*_*near*_ = 5 cm and *Z*_*far*_ = 150 cm. *s*_*W*_ and *s*_*H*_ are the width and height of the screen, respectively, and *s*_*W*_ = 105.2 cm, *s*_*H*_ = 59.2 cm.

The resultant coordinates are 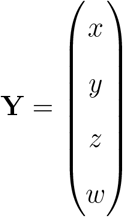.

Next, we can obtain the normalized screen coordinates, **N**:

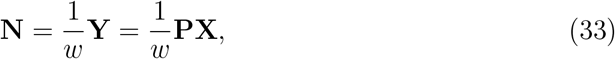

and *w* = −*Z*.

#### Object and object motion

Now consider an object whose location on the screen is defined as (*x*_0_, *y*_0_). Normalizing its screen coordinates by screen size, we have:

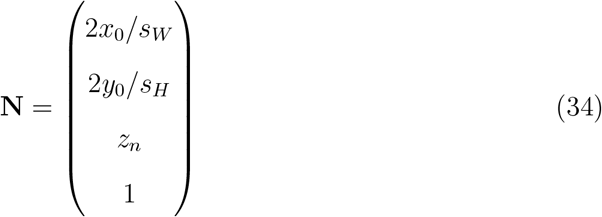

Object motion in the world is essentially a linear transformation, **M**, on the 3D coordinates, **X**. Consider only translation along three axes, 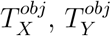, and 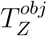:

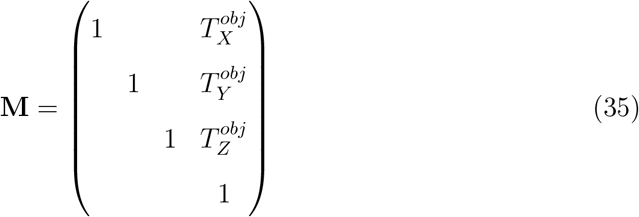

#### Self-motion

Similarly, self-motion is another linear transformation, **V**, on 3D coordinates. In this study, we only consider the rotation of the eye (i.e., rotation *θ* about the y-axis) and translation of the eye/head on a horizontal plane (i.e., translation along x-and z-axes, 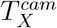 and 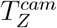):

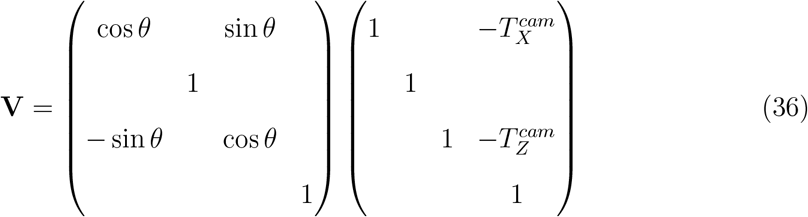

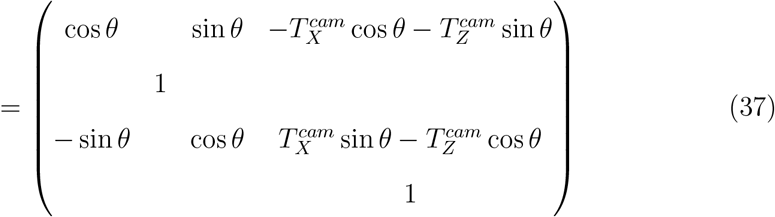

#### Retinal motion

Retinal motion is defined as a translation on the screen, with a certain amplitude, *l*, and direction, *α*. Therefore, the desired linear transformation, **A**, in screen coordinates is:

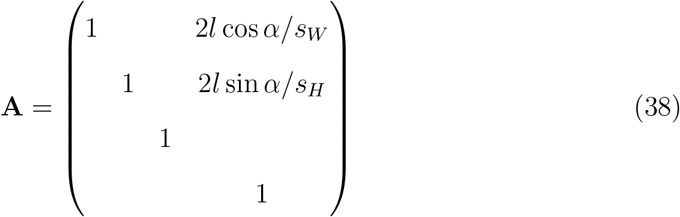

To present the desired retinal motion, **A**, on the screen, we need to find the correct 3D coordinates, **X**, object motion, **M**, and self-motion, **V**, such that:

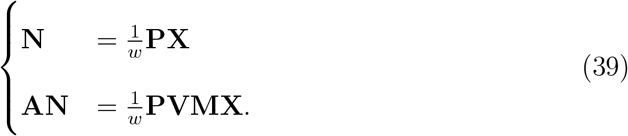

Here, the first equation specifies a mapping from world coordinates to normalized screen coordinates at the beginning of the trial *t*_0_, the second equation specifies such a mapping at a later time point *t*_0_ + ∆*t*, **P** is constant across time, **V** and **M** are the self-motion and object motion during time interval ∆*t*, respectively. Therefore, by solving the equations we can make sure that both viewing geometries produce the desired retinal motion.

#### Solution for the R viewing geometry

In the R geometry, the observer’s eye rotates around the y-axis and does not translate; therefore 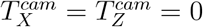. The initial z-coordinate of the object, *Z*, is the viewing distance, *Z* = −*f* . Solving for Equation (39), we have:

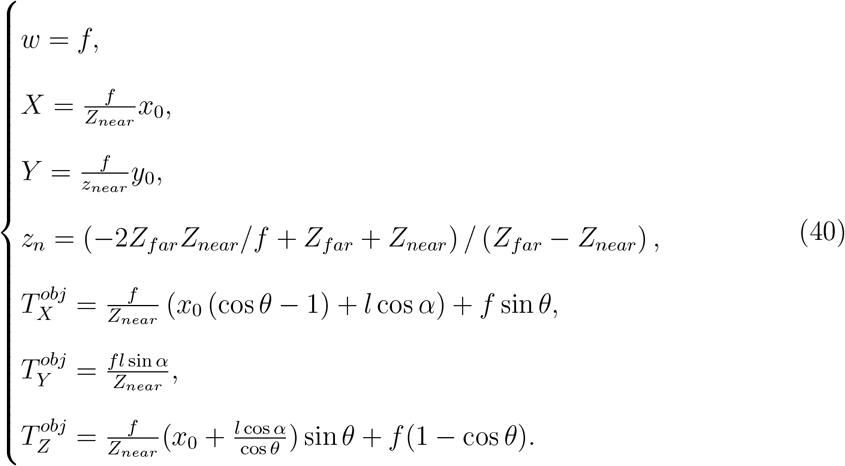

#### Solution of the R+T viewing geometry

In the R+T geometry, the object is located at a known depth, *Z*, and the observer’s eye translates along the x-axis while counter-rotating about the y-axis to maintain fixation, therefore 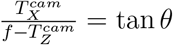. The object only moves along the y-axis, thus 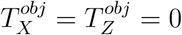. Solving for Equation (39):

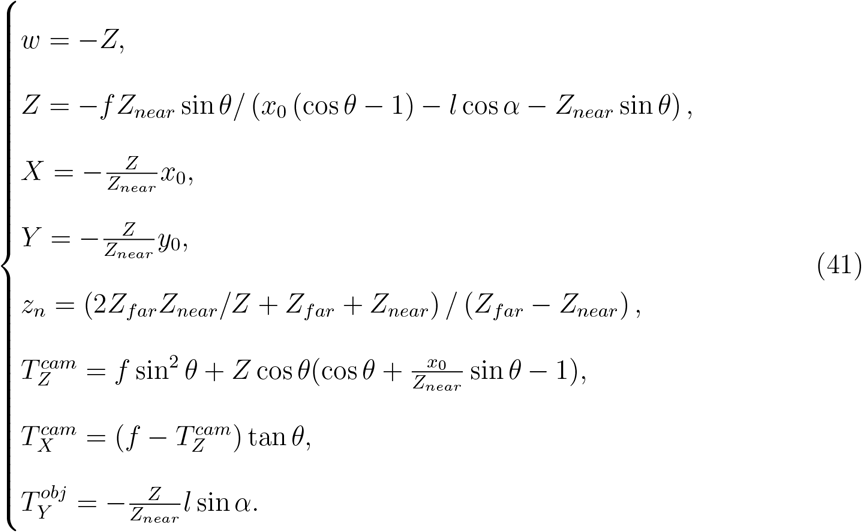

